# Genetic regulation of nascent RNA maturation revealed by direct RNA nanopore sequencing

**DOI:** 10.1101/2024.08.29.610338

**Authors:** Karine Choquet, Louis-Philippe Chaumont, Simon Bache, Autum R. Baxter-Koenigs, L. Stirling Churchman

## Abstract

Quantitative trait loci analyses have revealed an important role for genetic variants in regulating alternative splicing (AS) and alternative cleavage and polyadenylation (APA) in humans. Yet, these studies are generally performed with mature mRNA, so they report on the outcome rather than the processes of RNA maturation and thus may overlook how variants directly modulate pre-mRNA processing. The order in which the many introns of a human gene are removed can substantially influence AS, while nascent RNA polyadenylation can affect RNA stability and decay. However, how splicing order and poly(A) tail length are regulated by genetic variation has never been explored. Here, we used direct RNA nanopore sequencing to investigate allele-specific pre-mRNA maturation in 12 human lymphoblastoid cell lines. We found frequent splicing order differences between alleles and uncovered significant single nucleotide polymorphism (SNP)-splicing order associations in 17 genes. This included SNPs located in or near splice sites as well as more distal intronic and exonic SNPs. Moreover, several genes showed allele-specific poly(A) tail lengths, many of which also had a skewed allelic abundance ratio. *HLA* class I transcripts, which encode proteins that play an essential role in antigen presentation, showed the most allele-specific splicing orders, which frequently co-occurred with allele-specific AS, APA or poly(A) tail length differences. Together, our results expose new layers of genetic regulation of pre-mRNA maturation and highlight the power of long-read RNA sequencing for allele-specific analyses.

## Introduction

Premature messenger RNAs (pre-mRNAs) undergo several maturation steps prior to their release from chromatin. Most human pre-mRNAs have many introns that are removed through splicing, which takes place either co-transcriptionally or soon after transcription (Tilgner et al. 2012; Pandya-Jones and Black 2009; Yeom et al. 2021). Cleavage of the 3′-end occurs co-transcriptionally, after RNA polymerase II has transcribed through the poly(A) cleavage site, and is rapidly followed by polyadenylation, which consists of the addition of 200-250 adenines to the 3′-end of pre-mRNAs (Nicholson and Pasquinelli 2019). Alternative splicing (AS) and alternative cleavage and polyadenylation (APA), which are observed in 95% and 70% of human genes, respectively, allow extensive diversification and tailoring of cell proteomes (Pan et al. 2008; Derti et al. 2012; Wang et al. 2008).

Population-wide and quantitative trait loci (QTL) analyses have revealed an important role for common genetic variants in regulating AS and APA (Li et al. 2016; Pickrell et al. 2010; Garrido-Martín et al. 2021; Mittleman et al. 2020; Lappalainen et al. 2013; Ferreira et al. 2016; Mariella et al. 2019). Importantly, these variants harbor as much disease risk as expression QTLs, for pathologies ranging from immune disorders to schizophrenia (Li et al. 2016; Garrido-Martín et al. 2021; Walker et al. 2019; Raj et al. 2018). Yet, because these studies are generally performed with mature mRNA, they report on the outcome rather than the process, and may overlook ways in which variants can modulate pre-mRNA maturation. Indeed, mapping of alternative polyadenylation QTLs (apaQTL) in nuclear RNA revealed apaQTLs that were absent in whole cell RNA, while it also allowed to distinguish between co- and post-transcriptional regulatory mechanisms for whole cell apaQTLs (Mittleman et al. 2020). This highlights the necessity to understand how genetic variants impact RNA processing during the production and maturation of pre-mRNAs, which could in turn help explain their potential role in disease susceptibility.

Introns play a crucial role in AS control, as they include numerous sequence elements called splicing enhancers or silencers, which are bound by RNA-binding proteins (RBP) that regulate AS (Zhang and Chasin 2004; Fairbrother and Chasin 2000; Barash et al. 2010; Blencowe 2012). As such, the order in which introns are removed may determine how long splicing regulatory elements remain within a transcript to exert their influence on AS. Early gene-specific studies (Takahara et al. 2002; Schwarze et al. 1999; Kessler et al. 1993) and recent transcriptome-wide short- or long-read sequencing studies (Kim et al. 2017; Zeng et al. 2022; Drexler et al. 2020; Sousa-Luís et al. 2021; Gohr et al. 2022) demonstrated that introns are not always removed in the order in which they appear in the nascent transcript. Our recent work extended these results by studying the post-transcriptional splicing order for three to six consecutive introns (Choquet et al. 2023) using long-read direct RNA nanopore sequencing (Garalde et al. 2018). We revealed that multi-intron splicing order is predetermined, meaning that only a small subset of the possible splicing orders is used, which contributes to maintaining splicing fidelity. Surprisingly, we found that multi-intron splicing order is largely conserved between several human cell types, consistent with the conclusions from another independent study (Gohr et al. 2023). These findings suggest that the determinants of splicing order are hardcoded in the genome (Gohr et al. 2023). Yet, we and others have found only moderate associations between splicing order and several genomic features (e.g. intron and exon lengths, GC content, splice site sequences), suggesting the involvement of additional features (Zeng et al. 2022; Gohr et al. 2023; Choquet et al. 2023; Drexler et al. 2020).

Direct nanopore RNA sequencing (dnRNA-seq) (Garalde et al. 2018) yields long reads that allow for the simultaneous detection of several introns in the same molecule, making it ideal for investigating splicing order (Choquet et al. 2023). In addition, splicing and 3′-end processing are reciprocally regulated through interactions between RBPs over the last exon (reviewed in (Kaida 2016), emphasizing the importance of simultaneously interrogating distinct pre-mRNA maturation steps. Indeed, long-read sequencing studies have shown direct coupling between AS and APA in *Drosophila* and humans (Zhang et al. 2023; Alfonso-Gonzalez et al. 2023; Hardwick et al. 2022). Furthermore, long reads are particularly advantageous for allele-specific transcriptome analyses (Workman et al. 2019; Glinos et al. 2022; Tilgner et al. 2014), enabling the detection of several heterozygous single nucleotide polymorphisms (SNPs) in the same read and the assignment of reads to each parental allele to uncover splicing patterns that are specific to each allele. In addition, as pre-mRNAs from both alleles share the same cellular environment and potential technical variations during sample preparation (Demirdjian et al. 2020), any RNA processing differences between the two parental alleles should result from genetic variation rather than sample-to-sample variation. Lastly, long reads capture the maturation of pre-mRNAs that were previously difficult to characterize with short-read sequencing, such as transcripts from the highly polymorphic *HLA* genes (Tilgner et al. 2014; Cole et al. 2020).

In this study, we used dnRNA-seq of chromatin-associated, polyadenylated RNA to investigate allele-specific post-transcriptional pre-mRNA maturation in 12 human lymphoblastoid cell lines (LCL). We found dozens of instances of allele-specific splicing orders and identified proximal and distal genetic variants associated with splicing order changes, demonstrating that splicing order is at least partially determined by nucleotide sequence. Moreover, we uncovered several genes with significant differences in poly(A) tail length between alleles, which also exhibited allele-specific RNA abundance on chromatin. Deeper analysis of the highly polymorphic *HLA* class I transcripts revealed frequent allele-specific splicing order and poly(A) tail lengths. Together, this study describes a rich long-read sequencing dataset that can be used as a resource to study numerous aspects of allele- and compartment-specific gene expression, and reveals novel associations between genetic variation and key layers of pre-mRNA maturation.

## Results

### Subcellular dnRNA-seq across 12 lymphoblastoid cell lines

To explore the impact of genetic variants on nascent RNA processing, we leveraged naturally occurring heterozygous variants in lymphoblastoid cell lines (LCLs) from different individuals. We selected 12 LCLs derived from Yoruba (YRI) individuals that were part of the 1000 Genomes project, and for which gene regulatory variations have already been extensively studied (Li et al. 2016; Pickrell et al. 2010; Lappalainen et al. 2013; Mittleman et al. 2020). We performed cellular fractionation to collect cytoplasmic and chromatin-associated RNA from each LCL, followed by dnRNA-seq of poly(A)-selected RNA on a MinION or PromethION 2 Solo nanopore sequencing device (Table S1). Overall, we obtained medians of 6.07 and 1.70 million reads per LCL for the chromatin and cytoplasm fractions, respectively, for a total of 73.5 and 20.6 million reads per fraction. Moreover, our chromatin and cytoplasmic RNA datasets were characterized by median read lengths of 1255 and 854 nucleotides, respectively (Table S1), consistent with the presence of longer pre-mRNAs in the chromatin fraction (Fig. S1A). As demonstrated previously (Choquet et al. 2023), chromatin-associated RNA was enriched for partially spliced reads compared to cytoplasmic RNA across samples (Fig. S1B).

Each dark or light blue arrow represents one read, with light blue reads highlighting alternative splicing of the orange exon. The alternative intron is shaded in orange and reads are sorted based on the excision status of this intron. Heterozygous SNPs in LCL GM19102 are shown as vertical bars below the gene, with dark pink bars representing previously identified sQTL SNPs. B) Schematic of splicing order computation for groups of three introns. Each intermediate isoform *k* at splicing levels 1 (one intron excised, *L1*) and 2 (two introns excised, *L2*) is depicted. The associated read counts c_k_ are used to calculate splicing order scores for each of the six possible orders. The DRIMSeq differential transcript usage is used to test for differential splicing order using individual replicates, while allelic splicing orders from merged replicates are compared using a chi-square contingency test and the Euclidean distance between orders. C) Volcano plot showing the results of allele-specific splicing order analysis from merged replicates. Each dot represents one intron group. Intron groups that showed a significant difference in the analyses with individual replicates and merged replicates are shown in orange. D) Type of splicing order difference as a function of the Euclidean distance between allelic splicing orders. The different categories are schematized on the right, with the number representing the order in which each intron is removed, and are defined in more detail in the Methods. The transcript in grey has a lower top splicing order score than the ones in black. E) Splicing order plots showing allele-specific splicing orders for two example intron groups. The thickness and opacity of the lines are proportional to the frequency at which each splicing order is used, with the top ranked order per intron group set to the maximum thickness and opacity. d indicates the Euclidean distance between alleles.

To enable allele-specific analyses, we used the software LORALs (Glinos et al. 2022), in which nanopore sequencing reads are mapped to two personalized reference genomes per LCL, corresponding to each parental allele, thus avoiding filtering out reads because of SNPs that differ from the reference genome (Fig. S1C). For each read, the allele of origin was identified using HapCUT2 (Edge et al. 2017). On average, we were able to assign 22.7% of mapped reads to their parental allele of origin on chromatin, while this proportion was lower (11.7%) in the cytoplasm. We identified known allele-specific alternative splicing (AS) events (GTEx Consortium 2020) (Fig. S1D-E), such as in the gene *DPP7* (Fig. 1A), confirming the validity of our approach. Thus, dnRNA-seq enables allele-specific analyses of nascent and mature mRNA.

**Figure 1.**
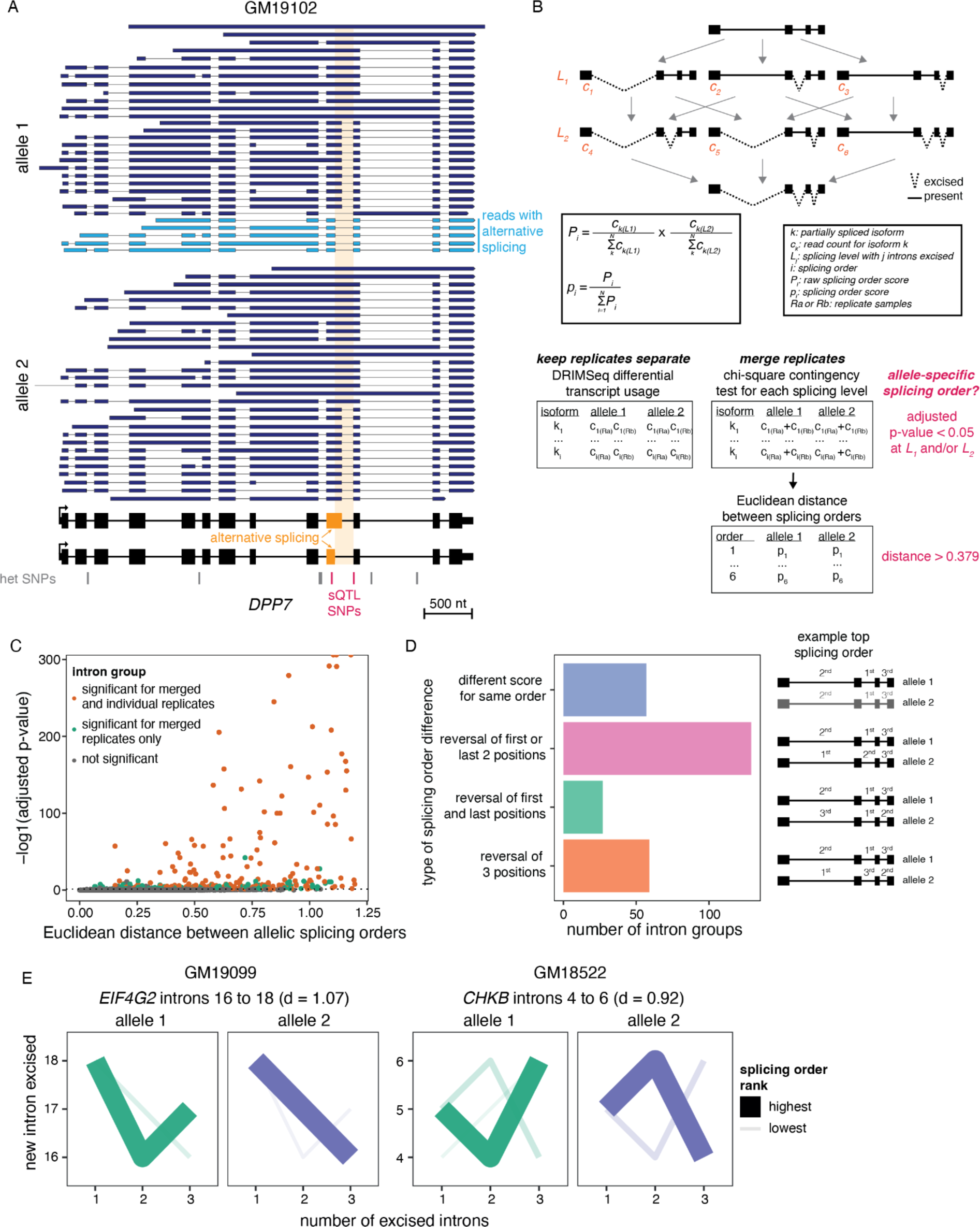
Subcellular dnRNA-seq reveals allele-specific splicing orders. A) Example of allele-specific splicing in *DPP7* in chromatin-associated RNA from LCL GM19102. 10% of reads mapping to each allele were randomly sampled. The gene structure is shown at the bottom, with exons as rectangles and introns as horizontal lines. The arrow represents the transcription start site.

### Splicing order variation between alleles

Previous work from our group and others suggest that splicing order is mostly determined by features that do not vary across cell types. We hypothesized that nucleotide sequence is the primary determinant of splicing order, in which case we would expect that some genetic variants lead to changes in splicing order. Thus, we sought to determine the extent of splicing order variation across individuals. We previously found good agreement between co- and post-transcriptional splicing order (Choquet et al. 2023). We focused on post-transcriptional splicing order because co-transcriptional splicing order is challenging to study with current dnRNA-seq read lengths, as most reads corresponding to elongating transcripts are completely unspliced (Drexler et al. 2020; Choquet et al. 2023). In addition, sQTL SNPs were found to be moderately enriched in introns that are removed post-transcriptionally (Garrido-Martín et al. 2021), making this intron population of particular interest. As in our previous study (Choquet et al. 2023), our analysis was limited to groups of three introns in highly expressed genes due to the current read length and coverage of dnRNA-seq (Fig. S1F). For each group of three proximal introns that met our coverage thresholds (see Methods) on both alleles, we extracted read counts for each isoform containing a combination of introns that had already been excised and introns that were still present (“intermediate isoforms”). We calculated their frequency relative to other intermediate isoforms with the same number of excised introns (“splicing level”). For each possible splicing order, the splicing order score was obtained by multiplying the frequency of the corresponding two intermediate isoforms (one per splicing level), as we recently described (Choquet et al. 2023) (Fig. 1B, Methods). We applied our strategy to each allele separately, yielding “allelic splicing orders” (Table S2). Similar to what we observed without allele specificity (Choquet et al. 2023), most intron groups displayed only 1-2 predominant splicing orders with scores > 0.25 on each allele, out of the six possible splicing orders for groups of three introns (Fig. S2A, Table S2).

We were able to compute allelic splicing orders for 58-1947 intron groups in each LCL (median of 638), depending on the sequencing depth and the number of heterozygous loci in highly expressed genes (Fig. S1F). We also assembled splicing orders without allele specificity (“allele-agnostic” splicing orders) for each LCL. Average allelic splicing order scores between alleles were highly correlated with allele-agnostic splicing order scores (Fig. S2B). The small number of outliers was mainly due to the absence or over-representation of some intermediate isoforms in the allelic splicing orders because of the presence of SNPs in only some introns of the group (Fig. S2C-D). Nevertheless, since the same SNPs are used to differentiate between alleles, this limitation did not prevent us from investigating splicing order differences between alleles.

Analysis of biological replicates from the same cell line, for which the RNA was collected and sequenced independently, or technical replicates, corresponding to different sequencing libraries from the same RNA sample (Table S1), showed strong reproducibility of allelic splicing orders, including for replicates sequenced with two versions of dnRNA-seq chemistries (SQK-RNA002 vs. SQK-RNA004) (Fig. S3A). Conversely, comparing splicing orders between alleles within each LCL revealed numerous intron groups with strong differences in splicing order scores (Fig. S3B). Of note, due to sequencing coverage constraints, allelic splicing order analysis was possible in only one LCL for almost half of intron groups (Fig. S4A). *HLA* class I genes were a notable exception, with all intron groups meeting our allelic analysis thresholds in five or more LCLs (Fig. S4B), likely due to their high density of polymorphisms (Voorter et al. 2016), thus enabling deeper insight into these genes.

To systematically identify splicing order differences between alleles across our dataset, for each intron group and LCL, we computed the Euclidean distance between vectors consisting of the splicing order scores for each allele (Fig. 1B). Intron groups that displayed the same splicing orders between alleles were characterized by a small distance, while differences in splicing order scores led to an increased distance (Fig. 1C, Fig. S4C). Applying this approach to our entire dataset, we found a median distance of 0.14, with a long right tail and a maximum distance of 1.20. Analysis of the Euclidean distance distribution between replicates of the same allele was used to define the threshold (interquartile range * 1.5 = 0.379) for identifying differences between alleles (see Methods) (Fig. S4C). As an orthogonal approach, we compared the distribution of intermediate isoform counts between alleles at each splicing level using a chi-square contingency test. Hereafter we refer to intron groups with a Euclidean distance > 0.379 and chi-square test FDR < 0.05 for at least one splicing level as displaying “allele-specific splicing order”.

More than half of intron groups displaying allele-specific splicing orders were detected in a single LCL (Fig. S4D), with those in *HLA* class I genes representing outliers that were detected in many LCLs (Fig. S4E). Of the 3564 unique intron groups that were analyzed, 159 (4.5%) showed differences in splicing order between alleles in at least one LCL (Fig. 1C). When counting all instances of intron groups with allele-specific orders even when they were observed in multiple LCLs, we detected 272 intron groups with allele-specific splicing order (Table S2). Intron groups that showed differences most frequently (57%) displayed changes in splicing order of 2 introns (e.g. *EIF4G2* and *CHKB*, Fig. 1E), while 22% showed changes in relative order of all three introns (Fig. 1D). To assess the reproducibility of allele-specific splicing orders, we analyzed differential intermediate isoform usage between alleles without merging replicates using DRIMSeq (Nowicka and Robinson 2016), a package developed for differential transcript usage analysis in small sample sizes (Fig. 1B). This revealed 62 unique intron groups with reproducible differences in differential intermediate isoform usage in at least one cell line. All of these overlapped with significant intron groups in the chi-square contingency test and 50 (81%) had a Euclidean distance above threshold with merged replicates (Fig. 1C). When counting all instances of intron groups even when they were detected in multiple LCLs, 143/144 and 110/144 overlapped with significant intron groups in the chi-square contingency test and had a Euclidean distance above threshold, respectively. Thus, although the sequencing depth does not allow to study allelic splicing order across multiple replicates for all intron groups, this analysis demonstrates the reproducibility of allele-specific splicing orders. Lastly, we investigated splicing order within 8292 unique intron pairs that did not meet our filters to be considered as part of groups of three introns, of which 114 showed significant allele-specific splicing order (Table S3). These findings indicate that splicing order is frequently modulated by allelic nucleotide sequence, solidifying our hypothesis that genetic sequence is the primary determinant of splicing order.

### Specific genetic variants are associated with allele-specific splicing orders

Considering our frequent observations of allele-specific splicing orders, we next wondered whether specific genetic variants were associated with splicing order variation. Most intron groups that showed allele-specific splicing orders contained numerous heterozygous SNPs that could underlie these differences. To identify specific SNPs associated with splicing order changes, we tested the association between each SNP and intermediate isoform counts across LCLs using the DRIMSeq transcript usage QTL (tuQTL) tool (Nowicka and Robinson 2016). SNPs within one gene that shared the same genotypes across samples were grouped into “haplotype blocks”. This revealed 472 unique SNPs in 127 haplotype blocks that were significantly associated with allele-specific splicing order of 29 intron groups in 17 genes, including seven *HLA* genes (Table S4). Previous work has revealed moderate associations between splicing order and several sequence features, including splice site strength and sequence content between the 3′ splice site and the branch point region (Zeng et al. 2022; Gohr et al. 2023; Choquet et al. 2023; Drexler et al. 2020). Consistently, a SNP at the last nucleotide (nt) of exon 8 in *CCM2*, which reduced the 5′SS strength of intron 8 from 8.81 to 2.69 (Yeo and Burge 2004), was associated with an excision order change of introns 6 to 8 (Fig. 2A). Moreover, a SNP in *CHKB*, located two nt from the top predicted branch point in intron 4 (Paggi and Bejerano 2018), was associated with an excision order reversal of introns 4 and 6, and a SNP located 7 nt from the 5′SS of *CYBA* intron 3 was associated with an excision order change of introns 3 to 5 (Fig. 2A).

**Figure 2.**
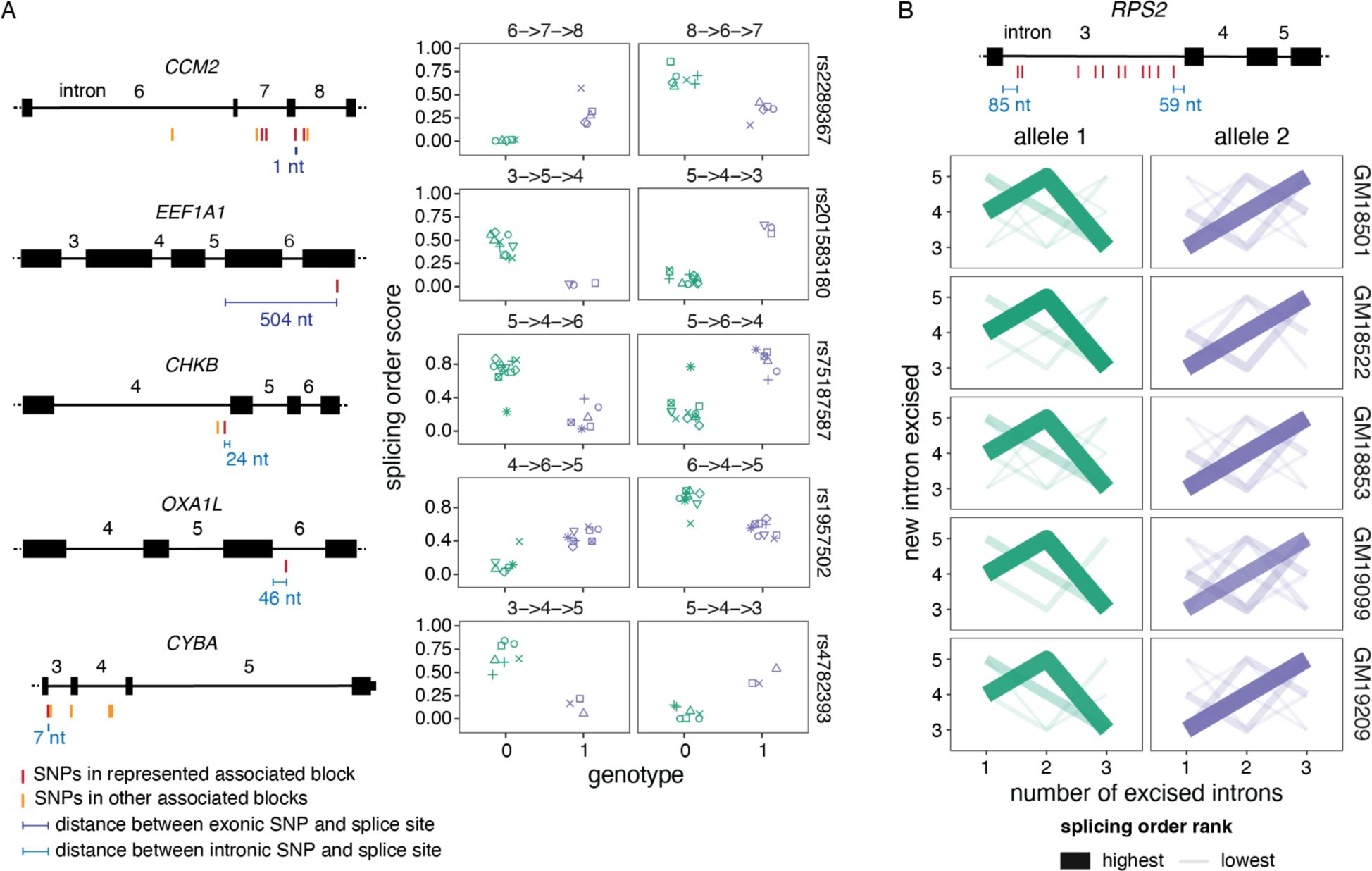
Splicing order changes are associated with specific genetic variants. A) Left: Schematic representation of each gene, with exons as rectangles and introns as horizontal lines. SNPs represented in the right plots and other SNPs in the same haplotype blocks are depicted in red, while SNPs that are significantly associated with splicing order of the same intron group but in a different haplotype block are shown in light orange. The distance between the SNPs represented in the right plots and the closest splice site are depicted in blue. Right: Examples of splicing orders in five genes that are significantly associated with the genotypes of the indicated SNPs. Each point represents one allele and the identity of the splicing order is shown at the top of each plot, with intron numbers separated by arrows. Each LCL is displayed in a different shape. B) Splicing order plot showing allele-specific splicing orders for introns 3 to 5 of *RPS2* in five LCLs. The thickness and opacity of the lines are proportional to the frequency at which each splicing order is used, with the top ranked order per intron group set to the maximum thickness and opacity. The gene structure is shown at the top, with exons as rectangles, introns as lines, and associated SNPs as in A).

Nevertheless, we also identified intron groups for which associated SNPs were outside of splice sites. In *OXA1L* introns 4 to 6, the only associated SNP within the intron group was 46 nt from the 5′SS. Moreover, *RPS2* intron 3 was removed prior to introns 4 and 5 on one allele, while this order was reversed on the second allele in five different LCLs (Fig. 2B). Multiple SNPs in linkage disequilibrium were associated with these *RPS2* allele-specific splicing orders (Fig. 2B). The majority were located within intron 3 but at least 59 nt from the splice sites, suggesting that some of these more distal intronic variants influence the timing of removal for this intron relative to its neighbors (Fig. 2B). Lastly, in *EEF1A1* introns 3 to 5, the closest associated SNP was located two exons and 504 nt downstream of the considered intron group (Fig. 2A), suggesting longer-range splicing regulation. Thus, our analysis suggests that splicing order may be regulated by changes to the consensus splice site sequences as well as more distal features, including outside of the affected exon.

We next explored potential genetic co-regulation of splicing order and AS. We found little overlap between SNPs associated with splicing order and GTEx sQTL SNPs (no common SNPs) or previously published sQTL SNPs identified in a cohort of YRI LCLs that includes the cell lines analyzed here (Li et al. 2016) (1 common SNP in the 3′UTR of *EEF1A1*, Table S4). Accordingly, comparison of transcript structure between alleles revealed few allele-specific AS events in intron groups with allele-specific splicing orders (6/272 LCL/intron group pairs, Table S2). Although the small number of intron groups studied here prevents us from drawing global conclusions, these findings hint that genetic regulation of splicing order is frequently distinct from that of AS in the analyzed genes. Nevertheless, we note that *RPS2* intron 3 displayed an alternative 5′SS on the same alleles that show delayed removal of this intron in three LCLs (Fig. 2B, Table S2), suggesting an interplay between AS and splicing order might occur in some genes.

### Most splicing orders lead to productive splicing and nuclear export

We next inquired whether all the observed splicing orders represent productive splicing paths, as some could lead to long-term intron retention and subsequent nuclear retention and decay. To determine whether allele-specific splicing orders impact the allelic balance of cytoplasmic isoforms, we compared allelic transcript ratios between chromatin and cytoplasm for all genes that showed allele-specific splicing order differences and for which alleles could be distinguished in the cytoplasm (n=92 gene/LCL pairs, “splicing order genes”). Globally, we found a significant correlation (R = 0.58) between chromatin and cytoplasmic allelic ratios for these splicing order genes (Fig. 3A), which was higher than the correlation observed when all genes detected in both compartments were considered (R = 0.37, Fig. S5A). The majority (76%) of gene/LCL pairs had an allelic ratio between 0.4 and 0.6 in both compartments (e.g. *GAPDH*, Fig. 3B), while 17% showed a skew in the same direction in both compartments or had a difference of less than 0.15 between compartments. Thus, only 7% of gene/LCL pairs showed a substantial difference in allelic ratios between compartments. For example, the most notable outlier was *RPS2*, for which we had detected strong splicing order differences (Fig. 2B). We observed that the allele associated with delayed removal of intron 3 accumulated on chromatin, while the allelic ratio was closer to 0.5 in the cytoplasm (Fig. 3C). Observation of the expected allelic ratio in the cytoplasm suggests that, for most cases, while removal of intron 3 is delayed on one allele, all transcripts are eventually spliced and exported to the cytoplasm. Moreover, the correlation between compartments for the splicing order genes increased (R = 0.72) when *RPS2* was removed. Thus, adopting different splicing orders does not appear to substantially influence relative mRNA abundance between alleles in the cytoplasm.

**Figure 3.**
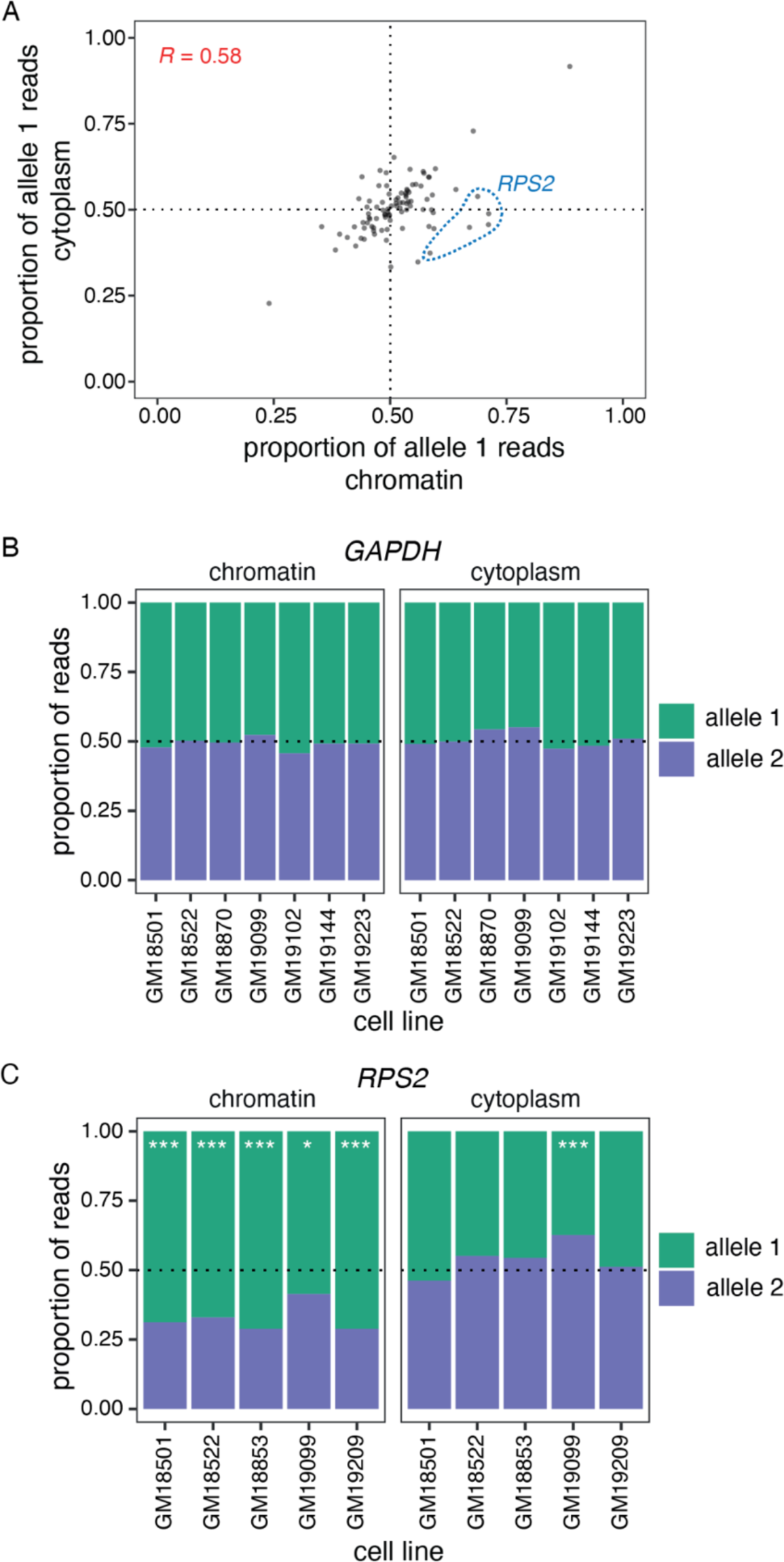
Splicing order changes do not alter cytoplasmic mRNA abundance. A) Correlation in allele-specific mRNA abundance between chromatin and cytoplasm for intron groups that showed significant allele-specific splicing orders. The proportion of allele 1 reads divided by the total number of reads for alleles 1 and 2 is shown for each subcellular compartment. Each dot represents one gene in one LCL. *RPS2* is highlighted in blue dotted circles for the five LCLs that showed splicing order changes in this gene and that are displayed in Fig. 2B. B) and C) Proportion of B) *GAPDH* (B) or C) *RPS2* reads mapping to each allele in chromatin and cytoplasm. The number of reads mapping to each allele on chromatin-associated or cytoplasmic RNA was compared using Qllelic or a two-sided binomial test, respectively. ***: p-value < 0.001 and proportion of allele 1 reads < 0.4 or > 0.6; *: p-value < 0.01 and proportion of allele 1 reads < 0.45 or > 0.55.

### Genetic regulation of nascent poly(A) tail length

Given the association between splicing order and genetic variation, we next asked whether other underexplored steps of pre-mRNA maturation are allele-specific. Poly(A) tails are added to newly synthesized pre-mRNAs on chromatin immediately after 3′-end cleavage. Following mRNA export to the cytoplasm, poly(A) tails are progressively shortened throughout the mRNA lifetime (Nicholson and Pasquinelli 2019). Nascent poly(A) tail length was long thought to be relatively constant across all nuclear pre-mRNAs (200-250 nucleotides) (Nicholson and Pasquinelli 2019), but we recently found substantial inter-gene variability of poly(A) tail length on chromatin-associated RNA (Ietswaart et al. 2024), which was closely associated with splicing progression (Choquet et al. 2023). However, how nascent poly(A) tail length is regulated or connected to splicing remains unknown. Therefore, we sought to establish whether poly(A) tail length is influenced by genetic variation. Poly(A) tail lengths were estimated from dnRNA-seq data and correlated well between replicates (Fig. S5B-C). As expected, poly(A) tails were longer and less variable on chromatin than in the cytoplasm, with respective medians of 176 and 97 nucleotides (Fig. S6A-B). We compared the distribution of poly(A) tail lengths between alleles in chromatin-associated and cytoplasmic RNA using a Wilcoxon rank-sum test. This showed 37 genes (65 gene/LCL pairs) with statistically significant differences in poly(A) tail length between alleles on chromatin-associated RNA, with only one gene, *HLA-DQA1,* identified as having a tail length difference in the cytoplasm (Table S5, Fig. S6C). This number is likely an underestimation, as poly(A) tail length measurements with dnRNA-seq are inherently noisy and require substantial coverage to detect differences. Indeed, 87% of genes with significant differences had at least 200 reads across both alleles, but this level of coverage was achieved by only 23% of assessed genes (Table S5). A correlation between allelic chromatin and cytoplasmic poly(A) tail lengths was not observed (Fig. S6D), consistent with the progressive deadenylation occurring in the cytoplasm (Nicholson and Pasquinelli 2019) and the positive correlation between the extent of deadenylation after nuclear export and cytoplasmic mRNA half-lives (Ietswaart et al. 2024). Among genes with allele-specific nascent poly(A) tail lengths, some showed robust differences across several LCLs. For example, in six LCLs, *ERAP2* displayed an average difference of 89 nt between the two alleles (average tail lengths of 233 and 143 nt per allele), while in five LCLs, *RPS2* exhibited an average difference of 39 nt (average tail lengths of 200 and 161 nt per allele) (Fig. 4A-B). Thus, our results suggest that nascent poly(A) tail length can be influenced by genetic variants, while such an association is rare for cytoplasmic mRNA.

**Figure 4.**
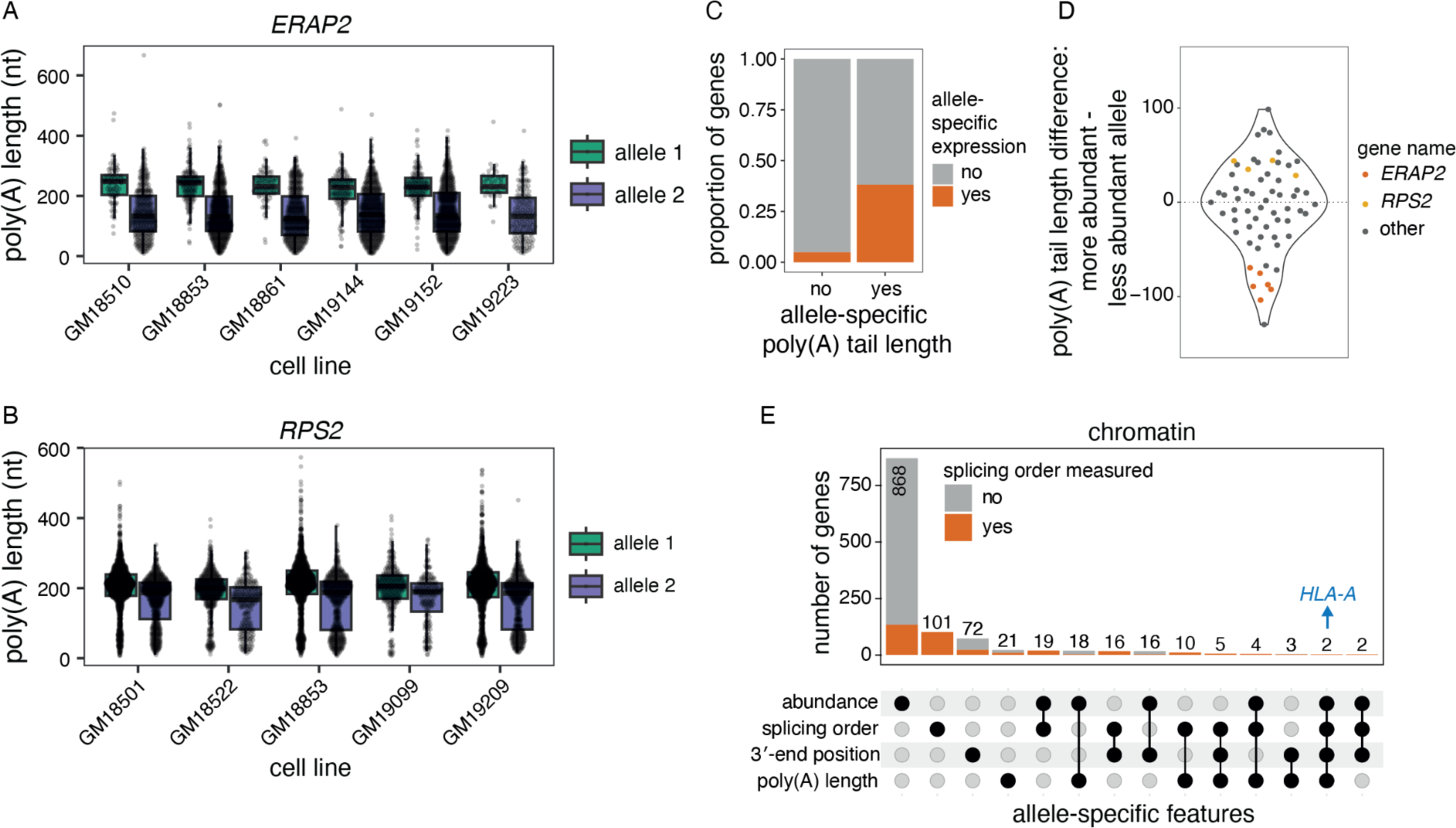
Genetic regulation of poly(A) tail length. A) and B) Examples of allele-specific poly(A) tail length in A) *ERAP2* and B) *RPS2*. Each dot represents one read. A two-sided Wilcoxon rank-sum test was used to compare tail length distributions between alleles for each LCL. All tests were statistically significant (adjusted p-value < 0.001). C) Proportion of genes that showed a skewed chromatin-associated RNA abundance ratio (allele 1 reads / total reads) out of genes with significantly different poly(A) tail lengths or not. The two groups were compared with a two-sided Fisher’s exact test (p-value = 9.232e-16). D) Violin plot of the distribution of the difference in poly(A) tail length between the more abundant and the less abundant allele for genes with a skewed chromatin-associated RNA abundance ratio. Each dot represents one gene in one LCL. E) UpSet plot showing the number of genes with skewed chromatin-associated RNA allelic abundance ratio (<0.4 or >0.6), allele-specific splicing order, significant differences in 3′-end position and/or significant differences in poly(A) tail length between alleles.

Hyperadenylation of nascent RNA has been associated with targeting for nuclear decay (Bresson and Conrad 2013; Bresson et al. 2015). Thus, we asked whether poly(A) tail length differences were accompanied by abundance imbalances between alleles. Genes with allele-specific poly(A) tail length were significantly enriched among genes with skewed nascent RNA abundance ratios (p-value = 9.232e-16, Fisher’s exact test, Fig. 4C). However, we found that the allele with the longest poly(A) tails could either be the least abundant (e.g. *ERAP2*) or the most abundant (e.g. *RPS2*) (Fig. 4A-B, D), suggesting that long poly(A) tails are not solely due to nuclear decay targeting and indicating a complex relationship between these two variables. Indeed, allele-specific abundance could also be the result of different RNA synthesis levels that may in turn impact poly(A) tail length.

Given the connections between 3′-end cleavage, polyadenylation and splicing (Kaida 2016), we investigated whether allele-specific poly(A) tail lengths, splicing order and/or alternative cleavage and polyadenylation (APA) co-occurred in some transcripts. We previously found an association between poly(A) tail length and splicing progression (Choquet et al. 2023). Thus, we first asked whether tail length differences on chromatin were primarily driven by partially spliced or fully spliced reads. The majority of genes showed significant differences in tail length for each read class (i.e. all reads, fully spliced reads only, partially spliced reads only) or only when all reads were considered, indicating that splicing progression is not the main reason for differences in poly(A) tail length. Nevertheless, we identified several genes that showed differences for one or two of these classes (Fig. S6E), such as *NOP56* (Fig. S6F). Overall, 21 gene/LCL pairs showed both allele-specific splicing order and poly(A) tail length differences (Fig. 4E), including *RPS2* (Fig. 4B). Future studies will be needed to establish whether those two features are independently regulated by genetic variants or if one is the cause of the other. We also asked whether allele-specific splicing order or poly(A) tail length are associated with APA. We extracted the genomic location of the 3′-end of each read and compared the distribution of 3′-ends between alleles. This approach recapitulated known allele-specific APA events, such as in *IRF5* (Fig. S6G) (Graham et al. 2007). Most genes with allele-specific APA did not show differences in splicing order or poly(A) tail length (n=88 gene/LCL pairs), while 10 and 25 gene/LCL pairs also displayed poly(A) tail length or splicing order differences, respectively (Fig. 4E, Table S6). Together, these results indicate that poly(A) tail length can be genetically regulated, and that this sometimes, but not necessarily, overlaps with other features such as pre-mRNA abundance, splicing progression, and APA.

### Multi-layered genetic regulation of HLA transcripts on chromatin

While many transcripts were heterozygous in a small number of LCLs, we reasoned that focusing on genes with higher level of variation across LCLs would provide greater power to dissect the influence of genetic variants on the different steps of pre-mRNA maturation and the interrelation between them. *HLA* genes, which are divided into class I (e.g. *HLA-A, -B, -C*) and class II (e.g. *HLA-DP, -DQ*, *-DR*) (Shiina et al. 2009), are the most polymorphic loci in humans (Voorter et al. 2016) and are highly expressed in LCLs (Fig. S7A). Accordingly, we detected *HLA* class I transcripts in most cell lines in our dataset (Fig. S4B, S4E). *HLA* genes encode immune receptors that play a central role in antigen presentation (Shiina et al. 2009). Given that alternative RNA processing can alter the identity or the levels of HLA proteins and affect their ability to present peptides to the immune system (Voorter et al. 2016), it is critical to understand how polymorphisms impact *HLA* pre-mRNA maturation. However, due to the technical challenges associated with analyzing *HLA* transcripts using short-read RNA-seq reads, *HLA* genetic variation has been primarily studied at the DNA level (Cole et al. 2020; Voorter et al. 2016), with little information as to whether and how this variation impacts pre-mRNA maturation. In all our analyses, *HLA* class I genes displayed significant differences between alleles. First, while the majority of intron groups with splicing order differences displayed only 1 or 2 different top splicing orders across LCLs (Fig. S4D), *HLA* class I genes frequently exhibited three or more orders (Fig. 5A), consistent with their higher degree of polymorphism. Furthermore, the majority of intron groups in these genes showed splicing order differences between alleles in most LCLs (Fig. 5A). To ensure that these were not due to technical artifacts when aligning partially spliced reads with short exons to the genome, we re-aligned reads to personalized *HLA* class I nascent transcriptomes (see Methods) and computed splicing order. This showed a very strong correlation (R > 0.99) between splicing orders calculated following genome or transcriptome alignment, irrespective of the proportion of alignment mismatches, which improved with updated dnRNA-seq chemistry (SQKRNA-002 vs. SQK-RNA004) (Fig. S7B-D). *HLA-B* exhibited one of the greatest Euclidean distances between splicing orders, with a maximum of 1.18 for introns 3-5 in LCL GM18501 and a median of 0.43 across intron groups and LCLs. Notably, six alleles (blue box in Fig. 5B) had strong usage of splicing orders 3->5->4 and 6->5->4, while they were never observed on other alleles. Splicing orders for both intron groups were associated with numerous SNPs, including some in introns 4 and 5 (Table S4). The other alleles displayed removal of intron 4 prior to intron 5, suggesting that some of these SNPs may be responsible for this strong splicing order reversal.

**Figure 5.**
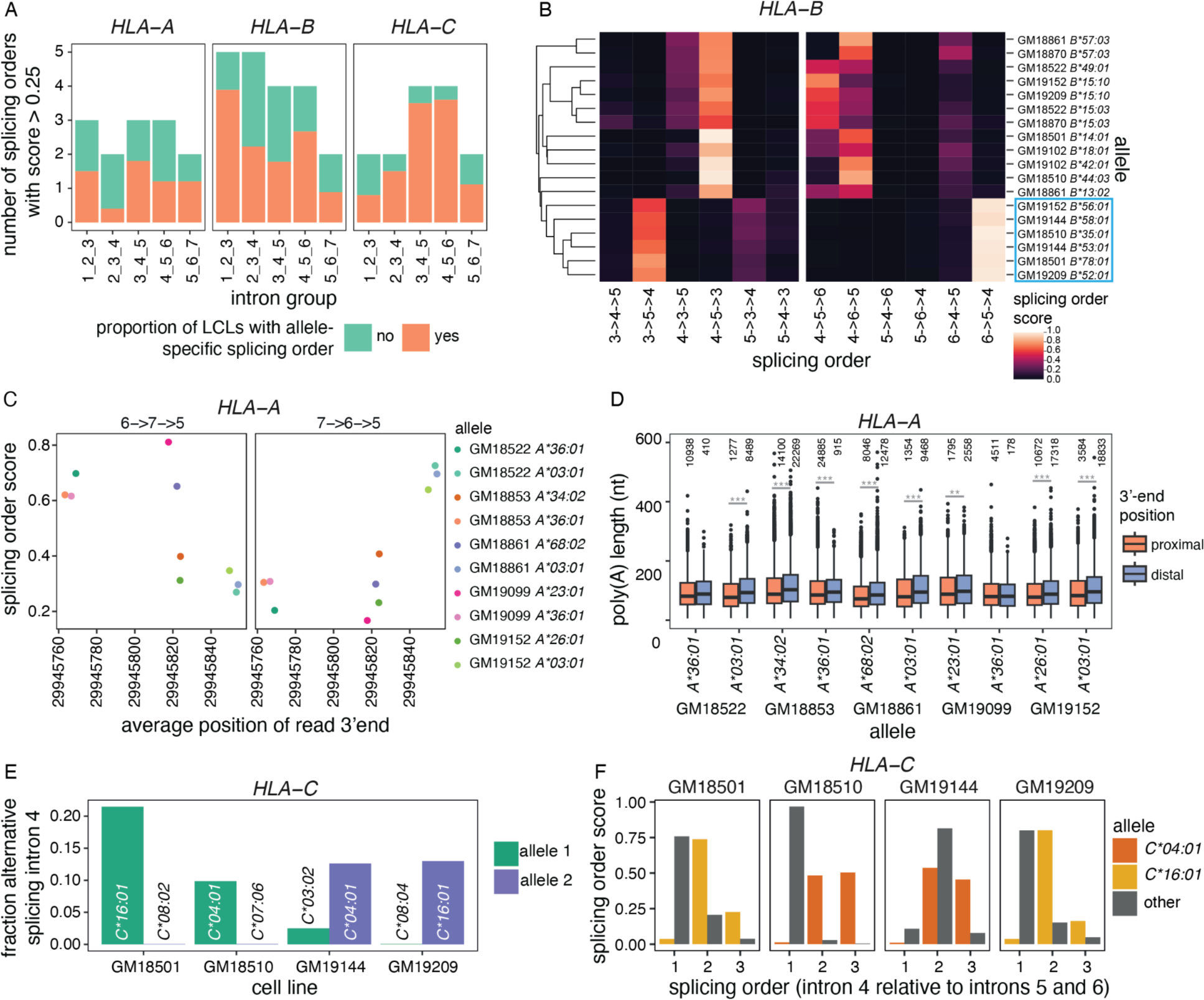
Multi-layered regulation of nascent *HLA* class I transcripts. A) Number of splicing orders observed across LCLs for each intron group in *HLA* class I genes. Each bar is colored based on the proportion of LCLs showing allele-specific splicing order for that intron group. B) Heatmap representing splicing orders for introns 3 to 5 and 4 to 6 in *HLA-B* across LCLs. Each line shows one allele and each column shows one possible splicing order. The squares are colored based on splicing order score. Alleles were clustered based on their genotypes across the *HLA-B* gene, so genetically similar alleles are located close to one another. The blue box highlights alleles that have delayed removal of intron 4 relative to other alleles. C) Splicing order scores as a function of average genomic read 3′-end position in *HLA-A*. Each dot represents one allele. The splicing order is shown at the top of each plot, with intron numbers separated by arrows. D) Poly(A) tail length distribution of *HLA-A* reads separated by allele, based on whether the read ends near the proximal or the distal 3′-end site. Poly(A) tail lengths were compared using a two-sided Wilcoxon rank-sum test. ***p-value < 0.001, **p-value < 0.01. E) Fraction of reads showing alternative excision of intron 4 in *HLA-C*, corresponding to exon 5 skipping. F) Splicing order score as a function of the position of intron 4 in each possible splicing order (removed first, second or third). The alleles for which there is AS of intron 4 in E) (*HLA-C*04:01* and *\*16:01*) show delayed removal of this intron relative to the other allele.

Interestingly, *HLA-A* was one of only two genes that displayed allele-specific splicing order, poly(A) tail length, 3′-end position, and abundance (Fig. 4E). Strikingly, these features appear to be co-regulated at the genetic level. Indeed, predominant usage of the proximal (e.g. *HLA-A*36:01*) or distal (e.g. *HLA-A*03:01*) 3′-end cleavage sites were respectively associated with strong usage of the splicing orders 6->7->5 or 7->6->5 (Fig. 5C). In addition, 3′-end cleavage at the distal site was associated with slightly longer poly(A) tails across most alleles (Fig. 5D), consistent with a previously reported positive correlation between 3′-UTR and poly(A) tail lengths (Alles et al. 2023). These observations suggest that 3′-end choice, poly(A) tail length, and excision order of the last two introns are coupled, although we cannot exclude that genetic variants independently act on each feature.

Additionally, we observed alternative splicing of exon 5 on *HLA-C*04:01* and *\*16:01* alleles in four LCLs (Fig. 5E), as previously reported (Ehlers et al. 2022). While the splicing orders varied between *HLA-C*04:01* and *\*16:01* alleles, they were all associated with delayed removal of flanking intron 4 (Fig. 5F), suggesting that AS and splicing order may be intimately linked in some genes. Despite allelic differences in pre-mRNA maturation, the allelic ratio was close to 0.5 for *HLA* class I genes across most LCLs, both on chromatin and in the cytoplasm (Fig. S7E), indicating that allele-specific splicing orders do not substantially impact mRNA abundance in these LCLs. Nevertheless, AS, APA and poly(A) tail length could impact translation, while splicing order may influence these aforementioned features, emphasizing the importance of understanding how the *HLA* transcript life cycle is regulated. Our analyses highlight that *HLA* class I genes can be regulated at these distinct levels, with potential downstream functional consequences.

## Discussion

Here, we performed dnRNA-seq of cytoplasmic and chromatin-associated, polyA-selected RNA in lymphoblastoid cell lines from 12 individuals for which multiple other functional genomics datasets already exist (Li et al. 2016; Mittleman et al. 2020; Pickrell et al. 2010; Lappalainen et al. 2013). To our knowledge, this is the first long-read subcellular RNA sequencing dataset across multiple individuals. In this study, we focused on the allele-specific analysis of two key aspects of pre-mRNA maturation, revealing that both proximal and distal variants are associated with splicing order changes and that nascent poly(A) tail lengths can differ by tens of nucleotides between alleles. Our findings highlight how analyzing specific RNA subpopulations can uncover new layers of genetic regulation during gene expression.

We observed that several groups of three introns displayed splicing order changes between the two alleles of each LCL. As both alleles were subjected to the same cellular environment and experimental procedures (Demirdjian et al. 2020), our findings demonstrate the crucial role of nucleotide sequence in splicing order determination and expand on previous studies showing that splice site and branch point sequences are moderately associated with splicing order (Zeng et al. 2022; Gohr et al. 2023; Choquet et al. 2023; Drexler et al. 2020). Consistently, several of the SNPs that we identified were located in splice sites and were associated with robust splicing order changes. Nevertheless, other SNPs were located further away from splice sites or from the intron groups themselves, though we note that short indels, which were not considered in this study, could also influence splicing order. Although future studies will be needed to confirm the causality of these genetic variants in modulating splicing order, our findings suggest that splicing order determination can involve regulatory elements located distally from intron-exon boundaries through mechanisms yet to be identified. Future work will need to be done to determine whether RBP binding events on nascent RNA contribute to splicing order determination. Genetic variants could also modulate splicing order by altering the secondary structure of pre-mRNA, which was shown to impact splicing efficiency and AS (Saldi et al. 2021; Taliaferro et al. 2016; Saha et al. 2020; McManus and Graveley 2011) through local or long-range intramolecular RNA-RNA interactions (Lovci et al. 2013; Kalmykova et al. 2021; Vorobeva et al. 2023; Margasyuk et al. 2023a). These long-range RNA interactions are enriched in introns compared to exons and most frequently occur through interaction of sequences within the same intron, which tend to be removed post-transcriptionally (Kalmykova et al. 2021; Margasyuk et al. 2023b, 2023a). Genetic variants, either individually or in combinations, could disrupt these structures to favor earlier or delayed removal of an intron relative to its neighbors, leading to the allele-specific splicing orders observed herein, especially when the associated SNPs are located away from splice sites (e.g. *RPS2*, *OXA1L*, *EEF1A1*). Future studies aimed at studying short- and long-range intramolecular RNA interactions in LCLs would help to shed light on such mechanisms.

Similar to our previous study on splicing order (Choquet et al. 2023), we focused on post-transcriptional splicing order, through which approximately 30-40% of mammalian introns are removed (Choquet et al. 2023; Yeom et al. 2021; Tilgner et al. 2012; Khodor et al. 2012). While our previous study suggested a high concordance between post- and co-transcriptional splicing orders (Choquet et al. 2023), post-transcriptional splicing order may provide additional time for genetic variants to influence splicing and may result in a higher occurrence of allele-specific splicing orders. As dnRNA-seq read lengths continue to improve, similar analyses of co-transcriptional splicing will help unravel the link between splicing timing and the effect of genetic variants. Furthermore, we were limited to analyzing 3,564 groups of three introns in highly expressed genes due to the current read length and coverage restrictions of dnRNA-seq (relative to the 119,936 intron groups in genes expressed in LCLs). Thus, the number of genes with allele-specific splicing orders or poly(A) tail lengths may not be reflective of the entire transcriptome and could be an under- or overestimation. In many intron groups, we did not find any genetic variant whose genotype was significantly associated with allele-specific splicing orders. While this is likely partly due to our small sample size, where allele-specific splicing orders were frequently observed in a single LCL, it is also possible that some allele-specific splicing orders result from combinatorial effects of several SNPs, as was shown for variants in DNA that physically interact to regulate gene expression (Corradin et al. 2016). This could be especially relevant if RNA structure is an important determinant of splicing order, and several genetic variants in a pre-mRNA favor the formation of alternative local and/or long-range interactions. Studies with larger cohorts will provide more power to identify instances consistent with such a mechanism.

We observed that splicing order changes did not affect cytoplasmic mRNA abundance, raising the question of the functional role of splicing order. We previously found that perturbing splicing order by blocking or mutating splice sites or depleting U2 snRNA severely impaired splicing fidelity (Choquet et al. 2023). However, most allele-specific splicing orders were not accompanied by increased AS or aberrant splicing. We propose that SNPs that impair splicing fidelity by modifying splicing order would be deleterious and have been negatively selected through evolution. Thus, common genetic variants such as the ones studied here are likely enriched for genes in which different splicing orders can be used without detrimental consequences or have advantages that led to their evolutionary selection. For example, in *HLA-C*, we found an association between exon 5 skipping and delayed intron 4 removal (Fig. 5E-F), consistent with our previous observations that introns flanking alternative exons tend to be removed later in motor neurons (Choquet et al. 2023). Future splicing order analyses of sQTL genes that did not have sufficient coverage to be investigated in this study could help to shed light on the relationship between AS and splicing order. Moreover, several recent studies have shown a connection between post-transcriptional splicing timing and response to stimuli (e.g. cell differentiation, neuronal stimulation, heat shock, etc.) (Yeom et al. 2021; Mazille et al. 2022; Mauger et al. 2016; Shalgi et al. 2014), where delayed splicing and nuclear transcript retention allow to form a pool of almost mature pre-mRNAs that can be spliced and exported upon specific signals. Some allele-specific splicing orders may provide a functional advantage for the response to stress or environmental signals that would only become apparent when performing a similar study in combination with different perturbations.

The use of dnRNA-seq enabled us to query allelic differences in poly(A) tail lengths of chromatin-associated RNA. In contrast to mature mRNAs, for which numerous studies have investigated the relationship between poly(A) tail length and mRNA stability or translation (reviewed in (Nicholson and Pasquinelli 2019)), the functional impact of shorter or longer poly(A) tail lengths globally is unclear, although hyperadenylation of some nuclear mRNAs contributes to their decay through the nuclear exosome (Bresson and Conrad 2013; Bresson et al. 2015). Some genes, where the least abundant allele had the longest tail (e.g. *ERAP2*, Fig. 4A), were consistent with this mechanism. On the other hand, genes for which the most abundant allele had the longest tail (e.g. *RPS2*, Fig. 4B) are more in line with our previous observations of a positive correlation between poly(A) tail length and the time that transcripts spend on chromatin (Ietswaart et al. 2024), suggesting that polyadenylation continues as long as the transcript is on chromatin. How polyadenylated mRNAs remain tethered to the chromatin is still unknown, but one hypothesis is that splicing completion acts as a licensing step for transcript release (Ietswaart et al. 2024; Brody et al. 2011; Hochberg-Laufer et al. 2019; Yeom et al. 2021). Although the individual contribution of these mechanisms to nascent poly(A) tail length control remains to be elucidated, our findings highlight previously underappreciated regulation at this level of gene expression.

*HLA* class I genes were not subjected to the sample size and heterozygosity limitations noted above, which allowed us to investigate the interplay between different pre-mRNA maturation steps in an allele-specific manner. Notably, we found an association between removal order of the last three introns and 3′-end APA in *HLA-A*, suggesting that the order or timing in which terminal introns are removed relative to their neighbors may be important for 3′-end choice, in addition to the known coupling between the splicing and 3′-end processing machineries (Kaida 2016). Further experiments are required to establish whether the co-occurrence of splicing order changes and APA is a special case in *HLA-A* or is widespread. Nonetheless, our results further emphasize the power of long-read sequencing to uncover connections between distinct pre-mRNA maturation steps and to decipher how genetic variants modulate these steps, which will help to unravel the mechanisms linking variants to human traits and diseases.

## Methods

### Cell lines

Human lymphoblastoid cell lines (LCLs) (Table S1) were purchased from the Coriell Institute for Medical Research. LCLs were maintained at 37°C and 5% CO_2_ in RPMI 1640 medium (ThermoFisher, 11875119) containing 10% FBS (ThermoFisher, 10437036), 100 U/mL penicillin, and 100 ug/mL streptomycin (ThermoFisher, 15140122). Cells were split every 3-5 days when they reached a density of approximately 1 million cells/ml.

### Cell fractionation and RNA extraction

Cellular fractionation was performed as described in steps 6 to 19 of (Drexler et al. 2021). For each LCL, 120-200 million cells were collected and centrifuged for 5 minutes at 500g, then split into aliquots of 10 million cells per cellular fractionation reaction (12-20 reactions total). At the end of the cellular fractionation, 3-4 chromatin pellets or 200 uL of cytoplasmic fraction were resuspended in 1 mL of Qiazol lysis reagent and stored at -80°C. For RNA extraction, samples were thawed for 2 min at 65°C, followed by addition of 200 uL of chloroform. Samples were vortexed for 15 seconds and centrifuged at 12,000g for 15 minutes at 4°C. The aqueous phase was transferred to a new tube, mixed with an equal volume of isopropanol, incubated at room temperature for 10 minutes and centrifuged at 12,000g for 10 minutes at 4°C. Pellets were washed twice with 1 mL of 75% ethanol, centrifuged at 7,500g for 5 minutes at 4°C and resuspended in nuclease-free water. Cell fractionations were verified by western blot as described in (Mayer and Churchman 2016) with antibodies against RNA polymerase II CTD phospho Ser2 (Active Motif, 61984) and GAPDH (ThermoFisher MA5-15738).

### Poly(A) selection and nanopore direct RNA sequencing

Poly(A)+ RNA was purified using the Dynabeads mRNA purification kit (ThermoFisher, 61006) according to manufacturer’s instructions, starting with up to 40 ug of chromatin-associated RNA and 75 ug of cytoplasmic RNA. Direct RNA library preparation was performed using the SQK-RNA002 or SQK-RNA004 kits (Oxford Nanopore Technologies) with 500-700 ng of poly(A)+ RNA according to manufacturer’s instructions with the following exceptions: the RCS was omitted and replaced with 0.5 uL water and the ligation of the reverse transcription adapter was performed for 15 minutes. For some initial samples (see Table S1), a yeast spike-in control was added to the libraries, as described in (Choquet et al. 2023). Sequencing was performed for up to 72 hours with FLO-MIN106D flow cells on a MinION or a PromethION 2 Solo device (Oxford Nanopore Technologies) (Table S1). For cell lines sequenced on a MinION, 3-6 flow cells were used on biological or technical replicates, while a single flow cell was used for each cell line sequenced on the PromethION 2 Solo.

### Nanopore data processing

Nanopore sequencing was performed with MinKNOW (release 22.03.5 or later). High accuracy basecalling was performed with Dorado v0.7.0 using default parameters and optional parameter - -estimate-poly-a. Basecalled BAM files from Dorado were converted to fastq using the samtools fastq command (Li et al. 2009). Reads were aligned to the reference human genome [ENSEMBL GRCh38 (release-86)] using minimap2 (Li 2018) with parameters -ax splice -uf -k14. As expected from the improved technology, libraries sequenced with SQK-RNA004 had a lower number of mismatches compared to those sequenced with SQK-RNA002 (Fig. S7B).

### Haplotype-aware alignment

Phased genotypes from LCLs were downloaded in VCF format from the International Genome Sample Resource (https://www.internationalgenome.org/data, GRCh38, release 20170504, downloaded in May 2019). We used the script process_vcf.sh from LORALS (Glinos et al. 2022) (https://github.com/LappalainenLab/lorals) to obtain per-individual VCF files that included only heterozygous SNPs. We then used the LORALS script make_new_vcf.sh and the initial alignments from chromatin-associated RNA samples (all replicates merged together for each cell line) to correct phased haplotypes in the VCF files and to generate two haplotype-specific reference genome fasta files per individual. The LORALS script hap_aligner.sh was then used to align reads (from chromatin-associated or cytoplasmic RNA) to these two genomes using minimap2 (Li 2018) and to select the highest scoring alignments as previously described (Glinos et al. 2022) to create a final BAM file from the two haplotype-aware alignments. The command extractHAIRS from HapCUT2 v1.3.3 (Edge et al. 2017) with options --nf 1 and --ont 1 and the script https://github.com/nanopore-wgs-consortium/NA12878/blob/master/nanopore-human-transcriptome/scripts/ase.py from (Workman et al. 2019) were used to assign reads to their alleles of origin, requiring that at least two heterozygous SNPs be present. Reads were classified as “not determined” if less than two informative variants were present or less than 75% of identified variants agreed with one of the two alleles.

### Determination of the excision status of introns

The excision status of introns was determined using our previously published method (Drexler et al. 2021). Reads that overlap introns from the hg38 RefSeq annotation were extracted using bedtools intersect (Quinlan and Hall 2010) with strand specificity. For each read/intron pair, we extracted the portion of the CIGAR string corresponding to the 50 nucleotides surrounding the 5′ and 3′ splice sites (SS) of the intron. Introns were classified as “excised” if the CIGAR string “N” (splicing event) started and ended within 50 nt of the 5′ and 3′SS and the size of this splicing event was within 10% of the annotated intron size. Introns were classified as “not excised” if there was no evidence of the CIGAR string “N”, and there was more than 50% coverage (CIGAR string “M”) in the 50 nt surrounding each splice site and more than 75% coverage within the region of intron that the read mapped to. Intron/read pairs where the read started within the intron (no coverage over the 5′SS) and met these criteria for the 3′SS were classified as “not excised”. Introns were classified as “skipped splice sites” if a splicing event (CIGAR string “N”) overlapped with greater than 50% of the 50 nt surrounding the 5′SS and/or the 3′SS and the portion of the intron within the splicing event was within 10% of the annotated intron size. These included alternative 5′ or 3′ SSs that are in an annotated exon and introns flanking skipped exons. Intron/read pairs that did not meet any of these criteria were classified as “undetermined”. Only introns with the splicing statuses “excised” or “not excised” were considered for splicing order analyses. To identify the proportion of partially spliced reads (Fig. S1B), all reads spanning two introns or more were considered. Reads that contained only “not excised” introns were considered “all unspliced”, reads that contained both “not excised” and “excised” introns were considered “partially spliced”, and reads that contained “excised introns” and no “not excised” introns were considered “all spliced”.

### Computation of splicing order

Splicing order for intron pairs was computed as described in (Drexler et al. 2020) using the excision statuses determined as outlined above. Reads spanning two consecutive introns with different excision statuses (one excised, one not excised) were extracted. The frequency of reads with the upstream intron removed or with the downstream intron removed was calculated for each allele. Intron pairs were used for downstream analyses if the number of reads per allele was higher than 10 and greater than twice the number of reads for which the allele could not be determined. Splicing order for groups of three introns was computed as described in (Choquet et al. 2023). Briefly, intron groups were analyzed when they met the following criteria: 1) each intron was retained in at least 10 reads in each allele; 2) each splicing level, defined as the number of excised introns within each read for the considered intron group, was represented by at least 10 reads that spanned all introns in the intron group of interest for each allele; 3) the number of reads per allele at each splicing level was greater than twice the number of reads for which the allele could not be determined. For duplicated intron groups with the same genomic coordinates within different transcripts, only one instance was kept. For overlapping intron groups in distinct transcripts that differ by an alternative splicing event (e.g. exon inclusion or exclusion), splicing order was measured separately for the two possible intron groups using different sets of reads that match each junction. For each splicing level *L* and allele A, the frequency *f_k_* of each possible intermediate isoform *k* was recorded by dividing the number of reads matching this intermediate isoform by the total number of reads at that splicing level. Next, we iterated through each level *L*, where for each observed intermediate isoform *k*, we identified the intermediate isoform(s) at the previous splicing level *L - 1* from which the isoform under consideration could originate. Those intermediate isoforms were connected within a possible splicing order path and their frequencies *f_k_* were recorded. After iterating through each level, the frequencies of patterns supporting each possible splicing order *i* were multiplied to yield the raw splicing order score *P_i_*, where *N* is the total number of intermediate isoforms supporting a given splicing order (4 for groups of 3 introns):

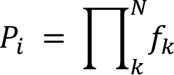

These raw scores *P_i_* were further divided by the sum of all raw scores for the considered intron group, where *n* is the total number of observed splicing orders for the intron group. This yielded the final splicing order score *p_i_* such that the sum of all scores *p_i_* was equal to 1:

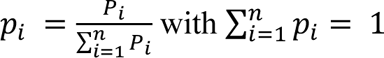

Only introns with “excised” or “not excised / present” statuses were considered for this analysis. Splicing order plots were produced in R with ggplot2 (https://ggplot2.tidyverse.org) with the following command:

~~~
ggplot(aes(x=splicing_level, y=new_intron_spliced)) + geom_line(aes(group=order_name, size=score, alpha=score))
~~~

Allele-agnostic splicing orders were computed in the same way, but using all reads (assigned to allele 1, allele 2, or undetermined).

### Identification of splicing order differences between alleles

To identify statistically significant differences in allelic splicing order for intron pairs, we compared the distribution of intermediate isoform (partially spliced) read counts between alleles using a two-sided Fisher’s exact test, followed by multiple testing correction with the Benjamini-Hochberg method. Intron pairs were considered to display allele-specific splicing when FDR < 0.05 and the absolute difference in the frequency of the upstream intron excised first between alleles was ≥ 0.1. To identify statistically significant differences in allelic splicing order for groups of three introns, we compared the distribution of intermediate isoform counts between alleles for each splicing level (one intron excised or two introns excised) using a chi-square contingency test, followed by multiple testing correction with the Benjamini-Hochberg method. To measure the extent of the difference between alleles, we calculated the Euclidean distance between splicing order scores by defining a vector with the six possible splicing order scores per intron group and allele, with the same order of splicing orders for the two alleles. We computed the Euclidean distance between the two vectors using the math.dist function in Python. To identify the proper threshold for splicing order differences between alleles, we computed the Euclidean distance for the same allele between biological or technical replicates of the same cell line for all intron groups that met coverage thresholds (N=914 intron group/LCL combinations). We calculated the interquartile range (IQR) of the combined distribution from all cell lines and defined the threshold for allele-specific splicing order as IQR * 1.5 = 0.379. Intron groups were considered to display allele-specific splicing when chi-square contingency test FDR < 0.05 for one or both splicing levels and Euclidean distance between allelic splicing order vectors > 0.379. As an orthogonal approach, we used the DRIMSeq (Nowicka and Robinson 2016) v1.30.0 differential transcript usage (DTU) workflow to identify allele-specific splicing orders without merging replicates. We extracted intermediate isoform counts from intron groups for which allelic splicing orders could be computed according to the filtering thresholds outlined in the “Computation of splicing order” section. Intermediate isoform counts were separated by splicing level (e.g. one intron excised or two introns excised), yielding three possible intermediate isoforms per splicing level. We used the script detect_differential_isoforms.py from rMATS-long v1.0.0 (https://github.com/Xinglab/rMATS-long) to execute the DRIMSeq DTU workflow, with each unique intron group at one splicing level representing a “gene” and each intermediate isoform representing a “transcript”. For each LCL, the two alleles represented the two groups being compared and the replicates were used as individual samples. Intron groups with adjusted p-value < 0.05 for at least one splicing level were considered to display allele-specific splicing order.

### Classification of splicing order differences

For each intron group with significant allele-specific splicing order, the top ranked splicing order was extracted for each allele and separated into individual introns. If two consecutive positions (first and second or second and third) were different between alleles while the remaining position (first or third) was the same (e.g. 1->2->3 vs. 2->1->3), the intron group was categorized as “reversal of first or last two positions”. If the first and last positions were different between alleles while the middle position was the same (e.g. 1->2->3 vs. 3->2->1), the intron group was categorized as “reversal of first and last positions”. If all three positions were different (e.g. 1->2->3 vs. 2->3->1), the intron group was categorized as “reversal of 3 positions”. If all three positions were the same, the intron group was categorized as “different score for same order”. The number of intron groups per category was counted and represented as in Fig. 1D.

### Testing for association between splicing orders and genetic variants

To identify SNPs associated with allele-specific splicing order, we used the transcript usage QTL (tuQTL) analysis workflow from DRIMSeq (Nowicka and Robinson 2016) with intermediate isoform counts from merged replicates. We extracted intron groups for which at least two LCLs displayed allele-specific splicing orders (chi-square contingency test p-value < 0.05 for one or both splicing levels and Euclidean distance > 0.379). We retrieved the genotypes for all SNPs in the corresponding genes, including 500 nt upstream and downstream, using the phased and corrected VCF files generated with LORALS (see above) and the window argument in the `dmSQTLdata` function. Each allele was considered to be a separate sample, with the genotype 0 for the reference allele and 2 for the alternative allele. Each unique intron group at one splicing level represented a “gene” and each intermediate isoform represented a “transcript”. Intron groups were filtered using the command `dmFilter,` requiring a minimum of ten alleles with at least 10 reads each and a minor allele frequency of at least 2 alleles. Within the tuQTL framework, SNPs within a gene that shared the same genotypes were grouped into a haplotype block (Nowicka and Robinson 2016). SNPs or haplotype blocks were considered to be significantly associated with splicing order if adjusted p-value < 0.05. For SNPs located in splice sites, the impact on splice site strength was assessed with MaxEnt (Yeo and Burge 2004). sQTL SNPs from (Li et al. 2016) were downloaded from http://eqtl.uchicago.edu/jointLCL/ on 2019/06/19 and SNP coordinates were converted from hg19 to hg38 using CrossMap (Zhao et al. 2014). sQTL SNPs identified in lymphoblastoid cell lines from GTEx v8 were obtained from GTEx_Analysis_v8_sQTL.tar, which was downloaded from https://www.gtexportal.org/home/downloads/adult-gtex/qtl. SNPs associated with splicing order and sQTL SNPs with the same coordinates were considered to overlap.

### Identification of allele-specific alternative splicing (AS) events

Using the chromatin-associated dnRNA-seq data from each LCL, two BAM files were created corresponding to reads mapping to each allele from the haplotype-aware alignment. Transcript isoforms were discovered and quantified from each BAM file using ESPRESSO (v1.4.0) with default settings and human genome reference and annotation [ENSEMBL GRCh38 (release-111)]. Transcripts with less than 10 reads on both alleles were excluded. Known and novel transcripts overlapping with genes with allele-specific splicing orders were extracted. For each pair of transcripts in each gene, the abundance between alleles was compared using a two-sided Fisher’s exact test. Multiple testing correction was performed using the Benjamini-Hochberg method. Transcripts were considered to have statistically significant allele-specific AS if the corrected p-value was < 0.05 and the odds ratio was < 0.5 or > 2. To identify the type of AS event between transcripts, we used the script classify_isoform_differences.py from rMATS-long v1.0.0 (https://github.com/Xinglab/rMATS-long) where the parameters –main-transcript-id and –second-transcript-id were the pairs of transcript isoforms considered to display allele-specific AS. AS categories “intron retention”, “alternative first exon”, “alternative last exon” and “complex” were excluded from further analyses. AS events in *HLA* genes were also excluded due to frequent genome alignment artifacts (see below) because of the small size of exon 6, leading to erroneous novel transcripts in ESPRESSO. For each intron group displaying allelic splicing orders, we defined a window corresponding to the start of the first intron -50 nt to the end of the last intron +50 nt and identified overlapping AS events using a custom Python script.

For comparing allele-specific splicing to known sQTLs, we used sQTLs identified in lymphoblastoid cell lines from GTEx v8, obtained from GTEx_Analysis_v8_sQTL.tar downloaded from https://www.gtexportal.org/home/downloads/adult-gtex/qtl. For each cell line, we identified sQTL SNPs that were heterozygous using the phased VCF files from LORALS. We extracted the corresponding sQTL introns and identified overlapping reads using bedtools intersect. For introns with more than 20 reads per allele, we compared alternative splicing between alleles using an intron-centric approach, as previously described (Choquet et al. 2023). Intron excision status was determined as described above. We compared the number of reads with “excised” vs. “skipped” or “undetermined” statuses on each allele using a two-sided Fisher’s exact test. Multiple testing correction was performed using the Benjamini-Hochberg method. The ratio of excised reads divided by the total number of reads (excluding “not excised” reads) was computed. Introns with an adjusted p-value < 0.05 and a difference between allelic ratios ≥ 0.1 were considered to display allele-specific alternative splicing. The same approach was used to analyze alternative splicing of *HLA-C*.

### Allele-specific transcript abundance

Reads were assigned to the genes they mapped to using bedtools intersect -s -F 0.5 -wo with the BAM file from haplotype-aware alignment and the hg38 RefSeq gene coordinates in BED format as input. The output file was merged with read to allele assignments from above and the number of reads per gene and allele was counted. Genes were included in this analysis if 1) they had more than 20 reads assigned to either allele, and 2) the number of reads assigned to each allele was at least two times higher than the number of undetermined reads. The allelic ratio was defined as the number of reads assigned to allele 1 divided by the number of reads assigned to alleles 1 or 2. We used Qllelic (Mendelevich et al. 2021) with 2 or 3 replicates per sample to identify genes with allelic imbalance in chromatin-associated RNA. We used a two-sided binomial test with p=0.5 to identify genes with allelic imbalance in cytoplasmic RNA due to the absence of replicates. For both approaches, multiple testing correction was performed with the Benjamini-Hochberg method. Genes with an adjusted p-value < 0.05 and an allelic ratio < 0.4 or > 0.6 were considered to have unbalanced or skewed RNA abundance. To assess the correlation in allelic ratio between subcellular compartments (Fig. 3A and S5A), only genes for which the ratio could be computed in both chromatin and cytoplasm were considered.

### Estimation and comparison of poly(A) tail lengths

Poly(A) tail lengths were estimated during basecalling with Dorado (see above) using the parameter --estimate-poly-a. We found that samples sequenced with SQK-RNA004 tended to have longer poly(A) tails than the corresponding replicates sequenced with SQK-RNA002, though poly(A) tail lengths remained highly correlated (Fig. S5B-C, S6C). Therefore, for cell lines sequenced with the two different chemistries, replicates were analyzed separately. Cell lines for which replicates were all sequenced with SQK-RNA002 on the MinION device had lower coverage and were therefore merged for further analyses. To compare poly(A) tail length between alleles, we filtered for genes that 1) had more than 20 reads assigned to each allele, and 2) the number of reads assigned to each allele was at least two times higher than the number of undetermined reads. Poly(A) tail length distributions were compared using a two-sided Wilcoxon rank-sum test. Multiple testing correction was performed with the Benjamini-Hochberg method. Genes with adjusted p-value < 0.05 were considered to have statistically significant differences in poly(A) tail length. For cell lines for which the replicates were not merged due to different chemistries being used, we required adjusted p-value < 0.05 in both replicates in order for the gene to have allele-specific poly(A) tail length. For comparing poly(A) tail lengths of partially spliced or fully spliced reads (as defined above), the same approach and thresholds were used.

### Comparison of 3′-end positions

The BAM file was converted to a BED file using BEDTools bam_to_bed (Quinlan and Hall 2010). For each read, the genomic position of the 3′-end was recorded in a strand-specific manner (end of the read on positive strand, start of the read on negative strand). The same coverage thresholds and treatment of replicates were used as for poly(A) tail length analysis. Distributions of 3′-end positions were compared using a two-sided Wilcoxon rank-sum test. Multiple testing correction was performed with the Benjamini-Hochberg method. Genes with adjusted p-value < 0.05 and at least 10 nt between the mean 3′-end position of each allele were considered to have statistically significant differences in 3′-end position (i.e. APA).

### Visualization

The coverage track in Fig. 1A was produced with pyGenomeTracks (Lopez-Delisle et al. 2021). The package ComplexUpset (https://github.com/krassowski/complex-upset) was used to represent the overlap of allele-specific features in each gene (Fig. 4E, S6E).

### Alignment to the HLA nascent transcriptome

In *HLA* class I genes, the short size of exon 6 led to artifacts suggesting skipping of this exon following alignment to the reference genome. Since we excluded any reads with exon skipping from splicing order analysis, we repeated splicing order computation with *HLA* class I genes after alignment to the transcriptome. For each *HLA* class I gene (*HLA-A*, *HLA-B*, *HLA-C*), we extracted the hg38 intron and exon annotations. Genomic coordinates were converted to coordinates from the start of each gene. We identified all possible combinations of retained introns, from 0 to 7 (completely unspliced), and re-constructed the nascent isoform sequences corresponding to each combination for each LCL using the haplotype-aware genome reference sequences generated above. From the haplotype-aware genome alignments, we extracted reads mapping to *HLA* class I loci and aligned them to the haplotype-aware *HLA* nascent transcriptomes using hap_aligner.sh as described above but with the minimap2 parameter -ax map-ont instead of -ax splice -uf -k14. For splicing order computation, the same strategy was used as above. For measuring the proportion of alignment matches per read in Fig. S7B, the CIGAR string was extracted with pysam (https://github.com/pysam-developers/pysam) (Li et al. 2009). The edit distance (NM tag) was subtracted from the total of matches (M) and insertions (I) and this number was divided by the total number of matches and insertions in the read (M + I - NM / M + I * 100).

### HLA typing

To identify the type of each *HLA-A, HLA-B* and *HLA-C* allele in our dataset, we used *HLA* typing data from (Abi-Rached et al. 2018) available through the 1000Genomes project FTP: ftp://ftp.1000genomes.ebi.ac.uk/vol1/ftp/data_collections/HLA_types/. After retrieving the *HLA* types for the 12 LCLs in our study, we extracted the sequences from the corresponding *HLA* alleles from the IPD-IMGT/HLA database (Robinson et al. 2015). For each LCL, we assembled a fasta file composed of the sequences of all possible variants of their assigned *HLA* alleles (hereafter “personalized *HLA* database”). We used bedtools (Quinlan and Hall 2010) getfasta to obtain the *HLA* sequences from the two haplotype-specific reference genomes for each LCL generated above from the dnRNA-seq data. These haplotype-specific *HLA* sequences were aligned to the corresponding personalized *HLA* database using minimap2 and the option --secondary=no. For each *HLA* class I gene, we used the primary alignment to assign each allele to one of the two known *HLA* types per individual.

## Competing interests

The authors declare no competing interests.

## Acknowledgements

We thank members of the Churchman lab for helpful discussions, advice, and assistance; Chantal Guegler, R. Stefan Isaac, Nicholas Kramer, Inés Patop and Ana-Maria Raicu for critical reading of the manuscript; and the Biopolymers facility at Harvard Medical School for sequencing services. This work was supported by the NIH (R01-GM136794, R21-HG011682 and R01-HG010538 to L.S.C.), the Fonds de Recherche du Québec - Santé, the Canadian Institutes of Health Research (post-doctoral fellowship awards to K.C.), Université de Sherbrooke and the Research Centre on Aging (start-up funds to K.C.).

## Author contributions

Conceptualization, K.C. and L.S.C.; Methodology, K.C. (lead) and L.S.C.; Investigation, K.C. (lead), L.-P.C., S.B., A.R.B.-K.; Software/Formal Analysis, K.C., L.-P.C. and S.B.; Writing – Original Draft, K.C. and L.S.C.; Writing – Review & Editing, K.C., L.-P.C., S.B., A.R.B.-K. and L.S.C.; Funding Acquisition, K.C. and L.S.C.; Supervision, K.C. and L.S.C.

## Notes

### Competing Interest Statement

The authors have declared no competing interest.

