## Supplementary Figures for "Genetic regulation of nascent RNA maturation revealed by direct RNA nanopore sequencing"

**Figure S1.** Quality control metrics of subcellular dnRNA-seq.

**Figure S2.** Characterization of allelic splicing orders.

**Figure S3.** Correlation in splicing order scores between replicates and between alleles.

**Figure S4.** Identification of allele-specific splicing orders.

**Figure S5.** Allele-specific mRNA abundance and poly(A) tail length quality control.

**Figure S6.** Allele-specific poly(A) tail length and 3′-end position.

**Figure S7.** Allele-specific analysis of *HLA* class I transcripts.

**Tables S1 to S6**, provided as a separate Excel file.

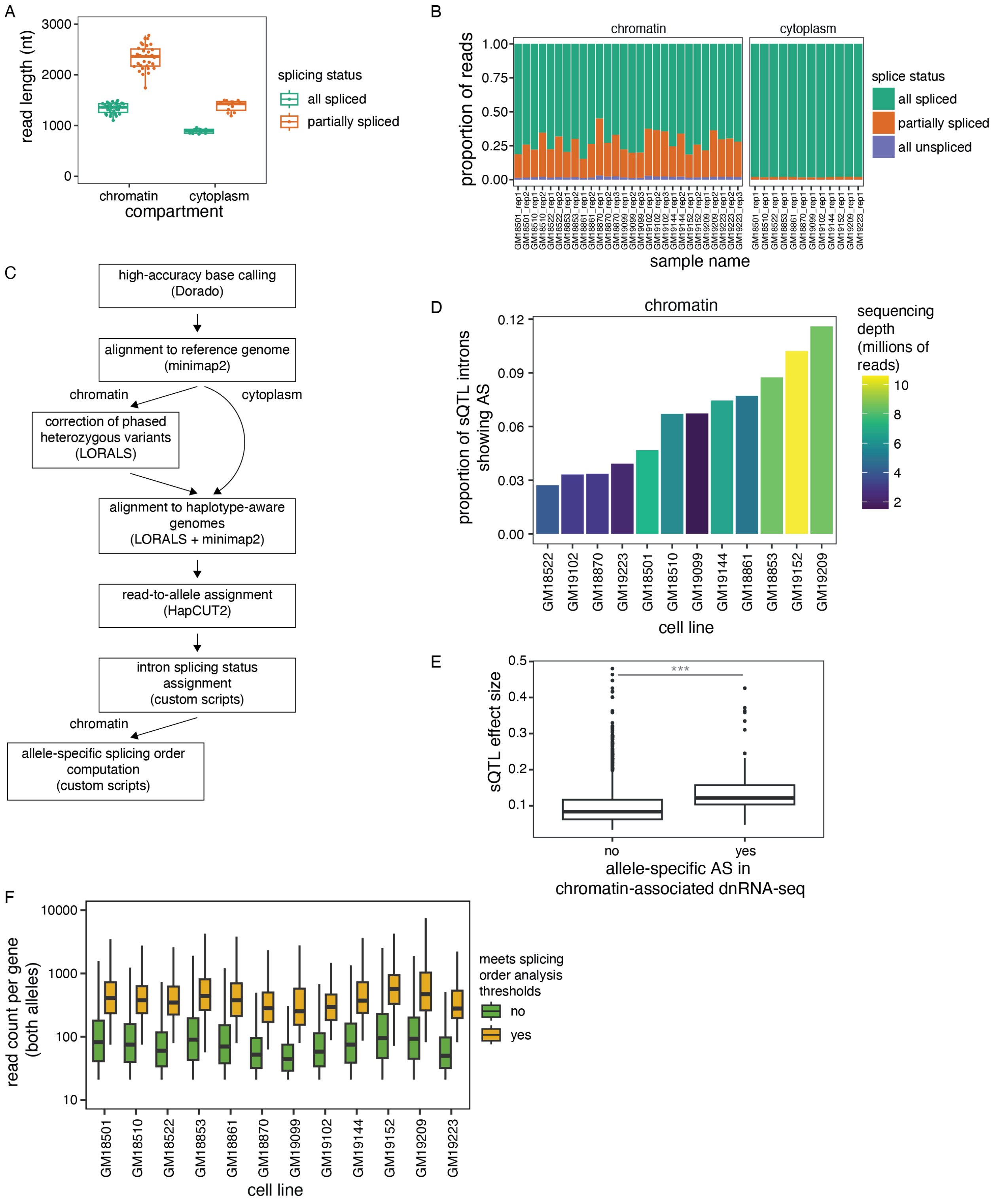


**Figure S1. Quality control metrics of subcellular dnRNA-seq (related to Figure 1).** A) Distribution of average read lengths per sample as a function of the read splicing status and the subcellular compartment. Each dot represents one sample. B) Proportion of reads spanning at least two introns that are all spliced, partially spliced, or all unspliced, in each subcellular compartment.

C) Workflow for the processing and allele-specific analysis of dnRNA-seq data. Correction of phased heterozygous variants is performed once per cell line using reads from chromatin-associated RNA, since they contain more introns and thus more information to determine phasing compared to cytoplasm. D) Proportion of GTEx sQTL introns showing alternative splicing (AS) in chromatin-associated dnRNA-seq. Bars are colored based on the total sequencing read depth for each cell line. E) GTEx sQTL effect size for introns that showed allele-specific AS or not in chromatin-associated dnRNA-seq. The two groups were compared using a two-sided Wilcoxon rank-sum test. ***: p-value $< 2.2e-16$. F) Distribution of the number of reads assigned to either allele for genes containing intron groups that meet the thresholds for splicing order analysis versus those that do not.

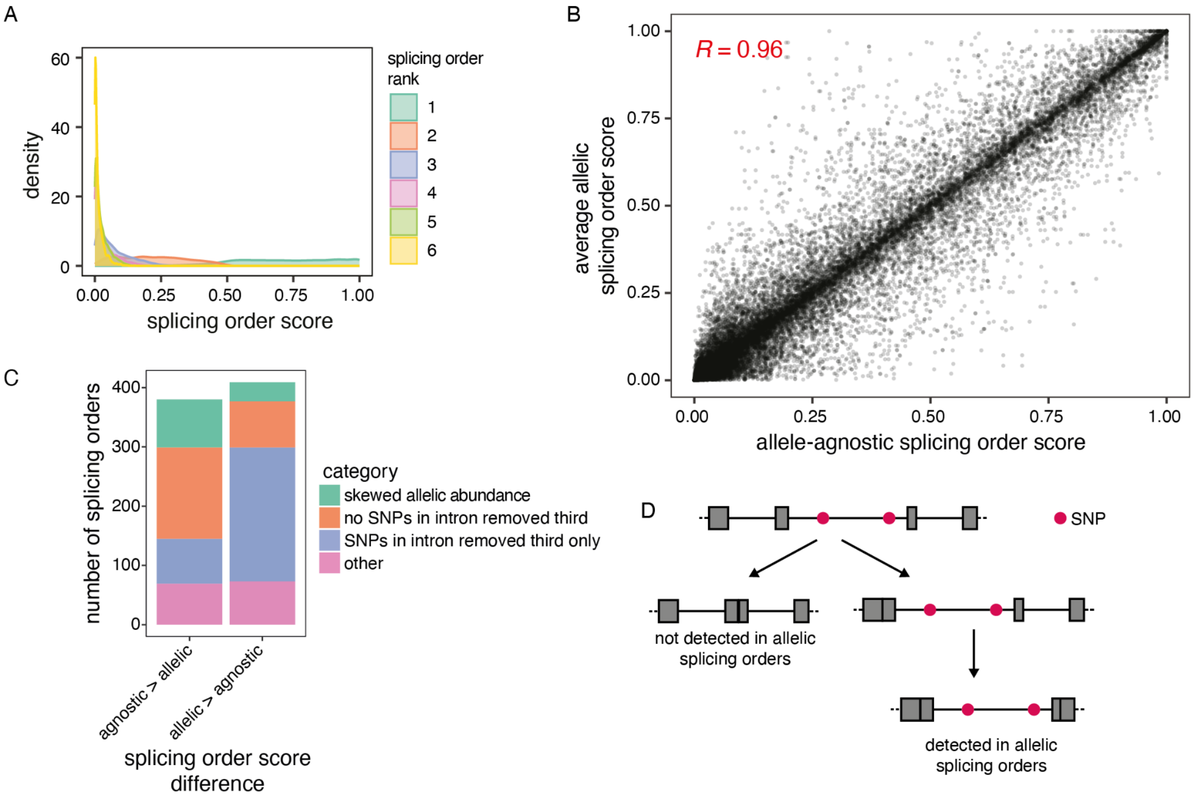


**Figure S2. Characterization of allelic splicing orders (related to Figure 1).** A) Distribution of splicing order scores across all LCLs and alleles, colored by splicing order rank. B) Correlation between allele-agnostic and average allelic splicing order scores across LCLs. Each dot represents one splicing order in one LCL. C) Classification of splicing orders based on whether there is a skewed abundance of pre-mRNA per allele (ratio of allele 1 reads / all reads < 0.4 or > 0.6) or whether there are SNPs in the intron that are removed third. Each observation refers to one splicing order in one cell line. Only splicing orders with a difference > 0.1 between allelic and allele-agnostic splicing order scores, and a score > 0.25 in at least one of the two approaches, are included. D) Schematic representing why some partially spliced reads are not detected (left) in allelic splicing orders, leading to a higher splicing order score for allele-agnostic splicing orders, or enriched (right) in allelic splicing orders, leading to a higher splicing order score for allelic splicing orders.

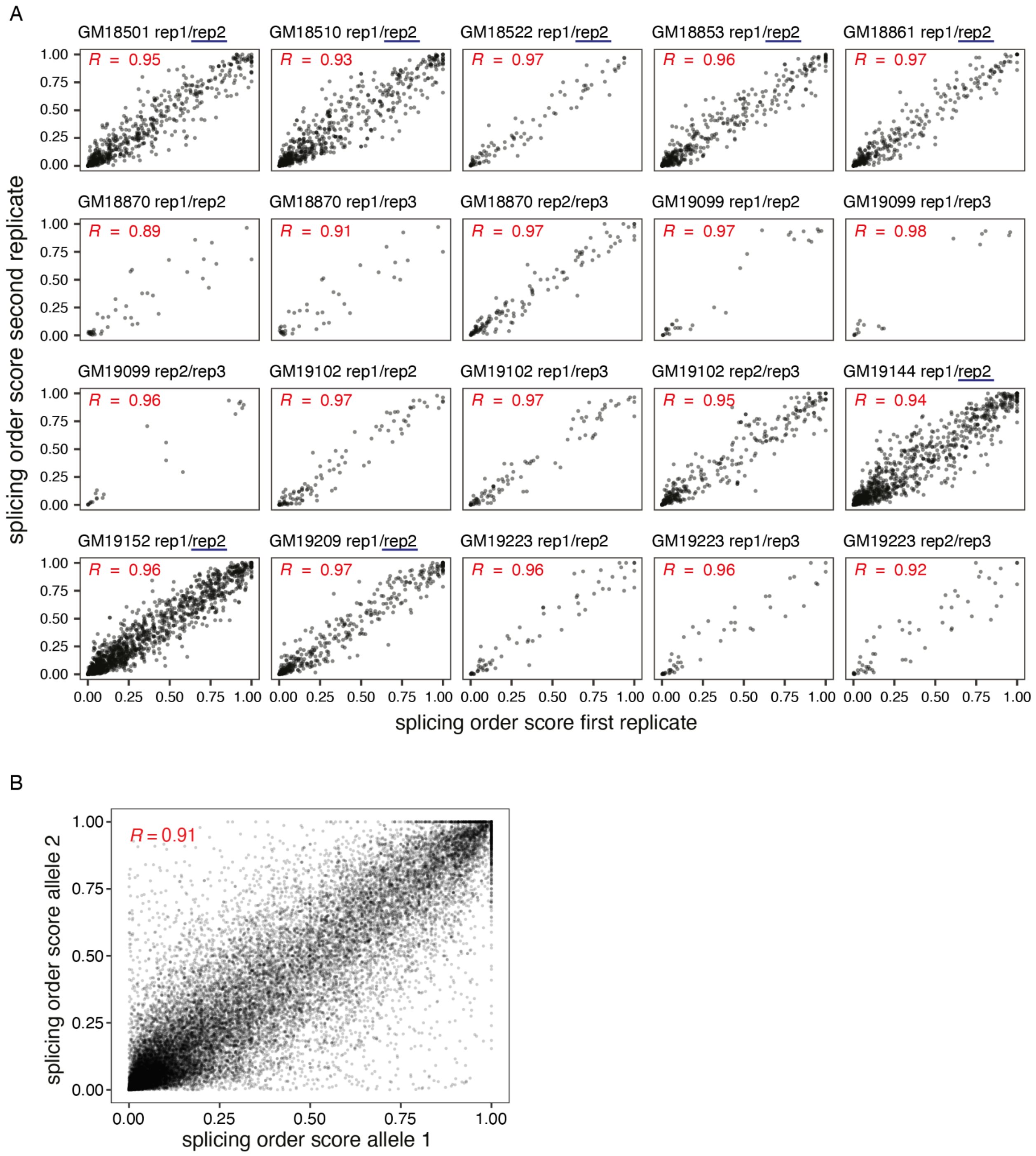


**Figure S3. Correlation in splicing order scores between replicates and between alleles (related to Figure 1).** A) Correlation in splicing order scores between biological or technical replicates of each LCL. Each dot represents one splicing order on one allele. Replicates sequenced with SQK-RNA004 are underlined in dark blue. For simplicity, "rep1", "rep2", and "rep3" refer to the first, second and third listed replicates in Table S1. B) Correlation in splicing order scores between alleles for all LCLs. Each dot represents one splicing order in one LCL.

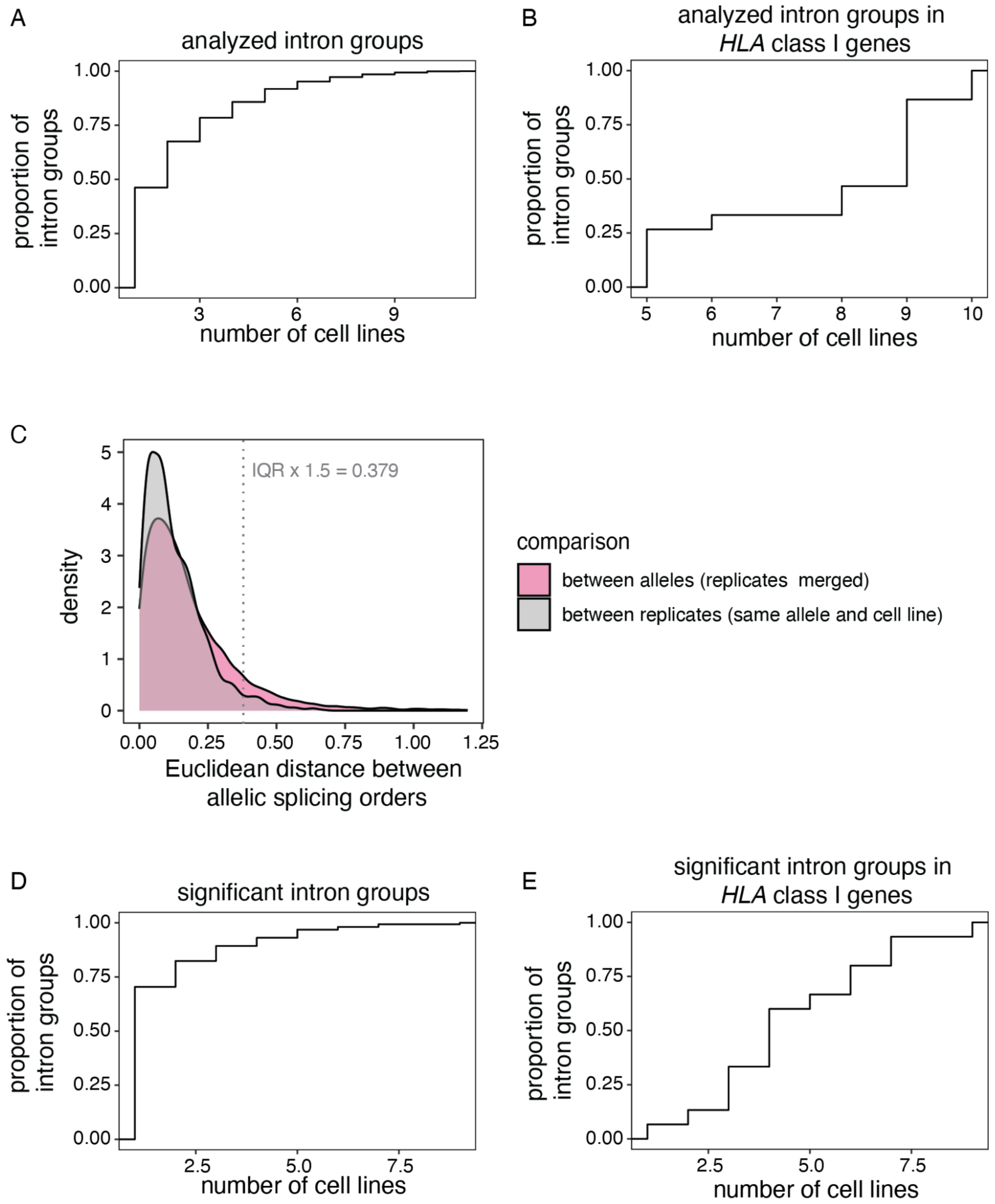


**Figure S4. Identification of allele-specific splicing orders (related to Figure 1).** A) Cumulative distribution frequency (CDF) of the number of cell lines in which each analyzed intron group could be detected in an allele-specific manner. B) Same as A), but only for intron groups in *HLA* class I genes. C) Distribution of the Euclidean distance between splicing order scores between replicates of the same allele (grey) or between alleles in merged replicates (pink) across all analyzed intron groups and LCLs. The threshold for a significant difference between alleles is shown as a grey dotted line (IQR: interquartile range of the distribution of splicing order scores between replicates).

D) CDF of the number of cell lines in which each analyzed intron group showed significant allele-specific splicing order. E) Same as C), but only for intron groups in *HLA* class I genes.

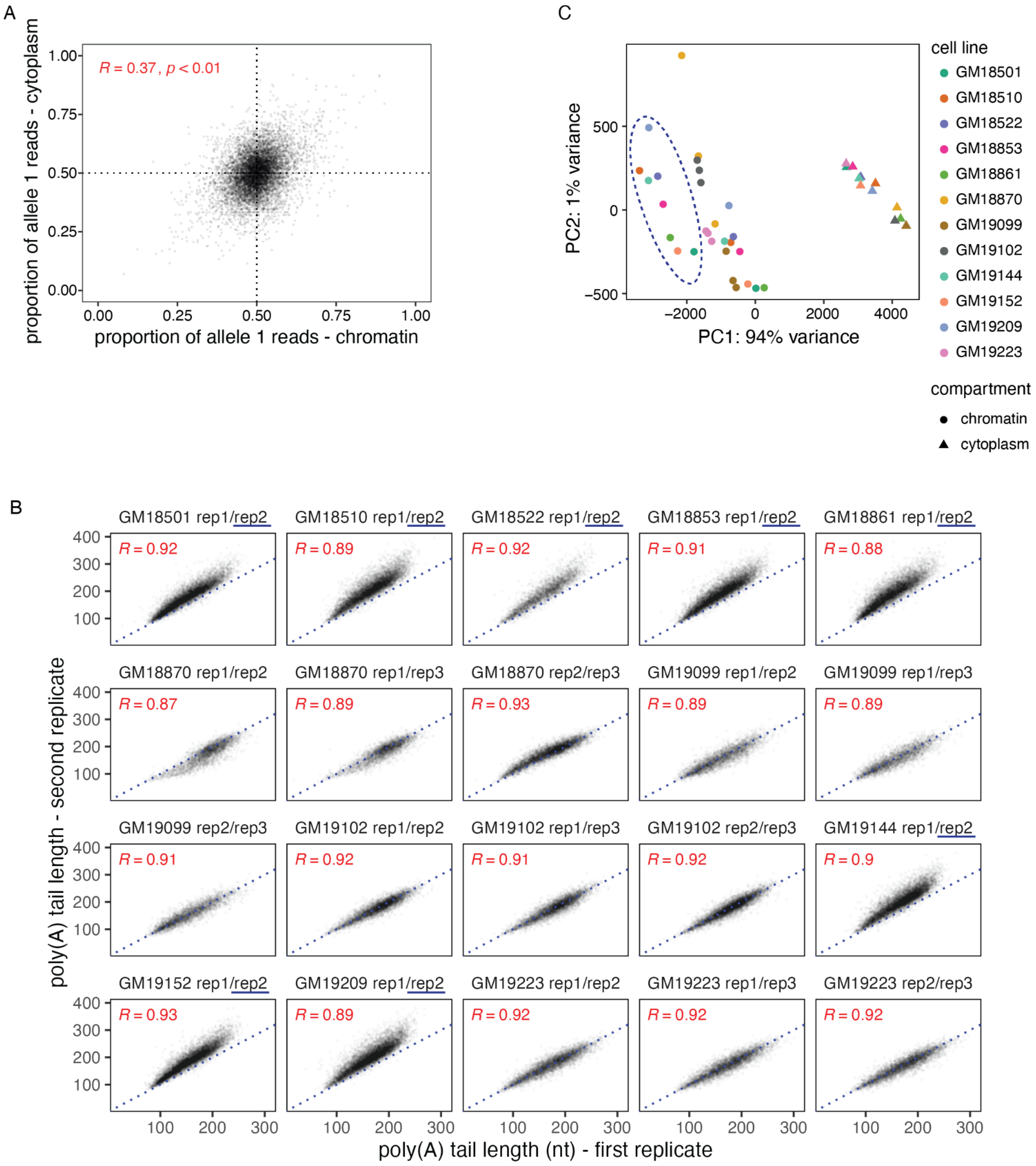


**Figure S5. Allele-specific mRNA abundance and poly(A) tail length quality control (related to Figures 3 and 4)**. A) Correlation in allele-specific mRNA abundance between chromatin and cytoplasm. The proportion of allele 1 reads divided by the total number of reads for alleles 1 and 2 is shown for each subcellular compartment. Each dot represents one gene in one LCL. All genes that met the coverage threshold in both compartments are shown. B) Correlation in chromatin-associated poly(A) tail lengths between biological or technical replicates. Each dot represents one gene. Replicates sequenced with SQK-RNA004 are underlined in dark blue. For simplicity, “rep1”, “rep2”, and “rep3” refer to the first, second and third listed replicates in Table S1. C) Principal component analysis of poly(A) tail lengths across replicates and subcellular compartments. Replicates sequenced with SQK-RNA004 are circled with a dotted dark blue line.

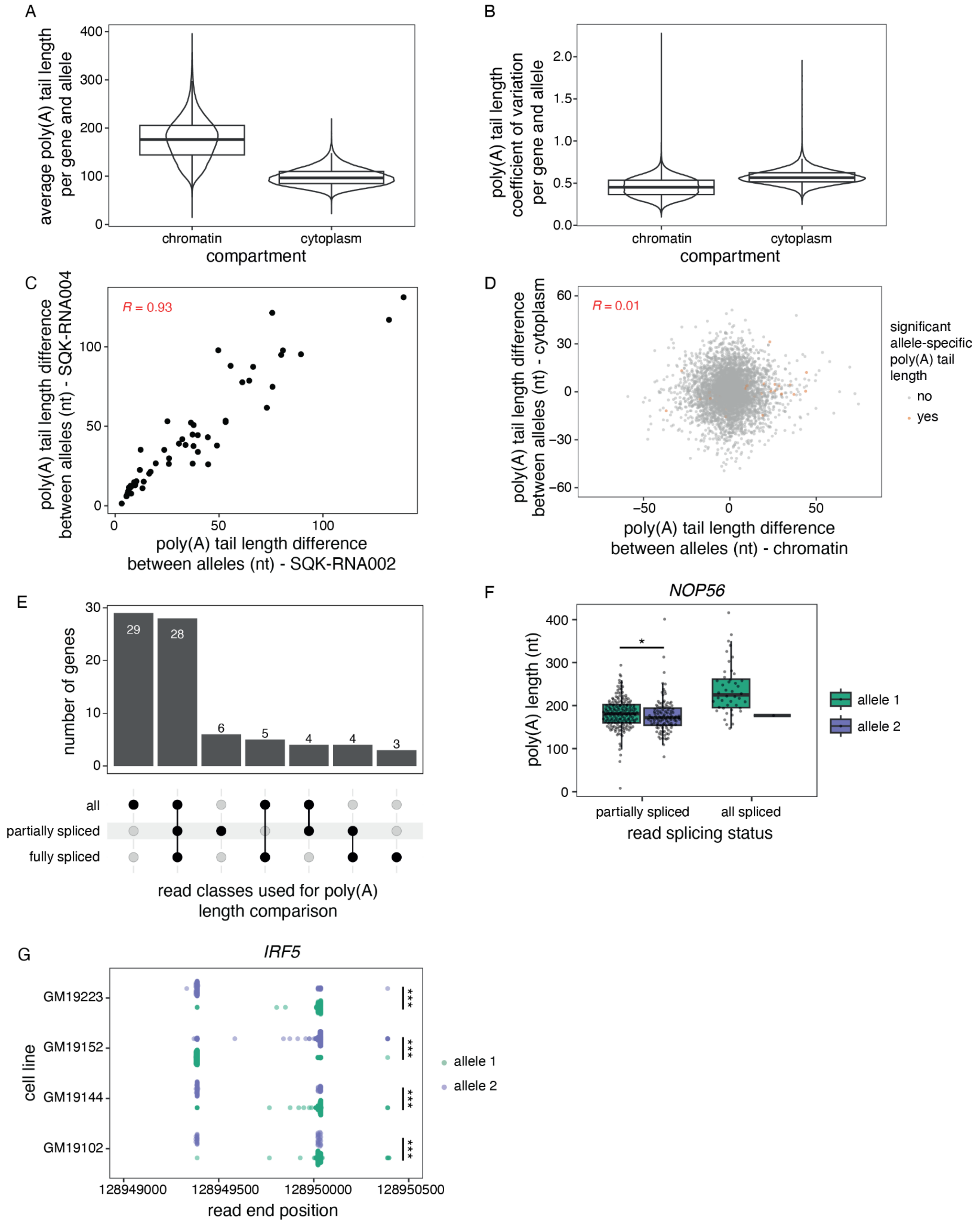


**Figure S6. Allele-specific poly(A) tail length and 3′-end position (related to Figure 4).** A) Distribution of the average poly(A) tail length per gene and allele across all LCLs for each compartment. B) Distribution of the poly(A) tail length coefficient of variation per gene and allele across all LCLs for each compartment. C) Correlation in the absolute poly(A) tail length difference between alleles for genes that showed a significant difference in chromatin-associated poly(A), in LCLs for which one technical replicate was sequenced with SQK-RNA002 and the other with SQK-RNA004. Each dot represents one LCL/gene pair. D) Correlation in poly(A) tail length difference between alleles (allele 1 - allele 2) between chromatin-associated and cytoplasmic RNA. Genes with significant differences in poly(A) tail length on chromatin are shown in orange. E) UpSet plot showing the number of genes with a significant difference in poly(A) tail length between alleles for different classes of reads (all reads, partially spliced reads and fully spliced reads). F) Poly(A) tail length distribution for partially spliced and all spliced reads from *NOP56* in GM19102. A two-sided Wilcoxon rank-sum test was used to compare tail length distributions between alleles for each read splicing status. *: adjusted p-value < 0.05. G) 3′-end position of reads mapping to *IRF5* in four LCLs. Each dot represents one read. The x-axis shows allele-specific differential accumulation of reads at two distinct positions. A two-sided Wilcoxon rank-sum test was used to compare 3′-end position distributions between alleles for each cell line. ***: p-value < 0.001.

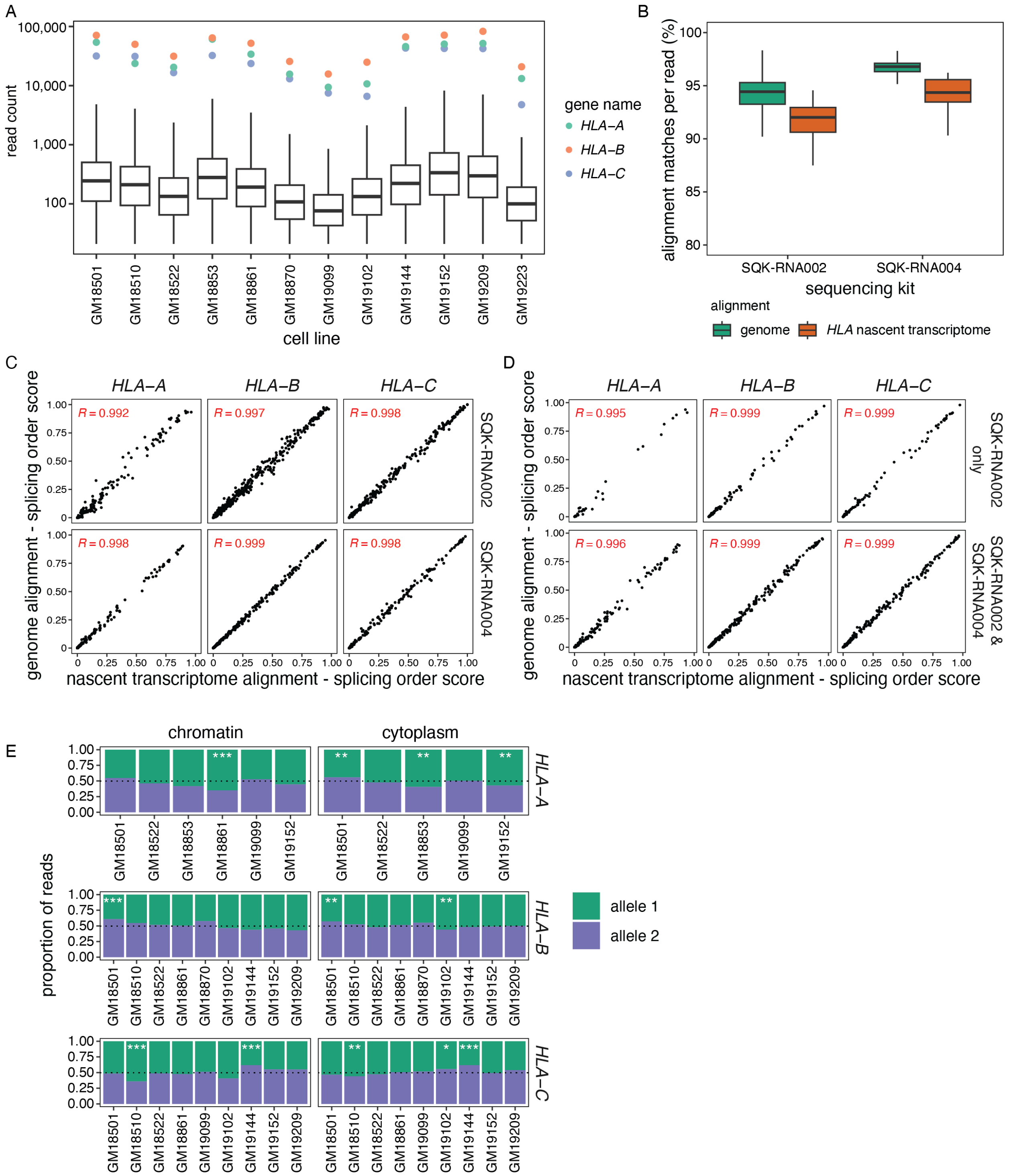


**Figure S7. Allele-specific analysis of *HLA* class I transcripts (related to Figure 5).** A) Boxplots representing the read count distribution of genes with at least 20 reads in each LCL. Colored dots indicate the read counts for *HLA* class I genes. B) Percent of nucleotides per read matching the reference sequence for alignment to the genome or to the *HLA* nascent transcriptome, as a function of the Oxford Nanopore Technologies sequencing chemistry used. C) Correlation in splicing order scores obtained from aligning reads to the genome or to the nascent transcriptome (see Methods) for individual replicates, separated according to the sequencing chemistry used. Each dot represents one splicing order on one allele. D) Same as C), but for splicing order scores obtained from merged replicates, separated based on whether all replicates were sequenced with SQK-RNA002 or with a combination of the two kits. E) Proportion of reads mapping to each allele of *HLA-A, HLA-B* and *HLA-C* in chromatin and cytoplasm. Only LCLs for which allele-specific RNA abundance met the required coverage thresholds are shown. The number of reads mapping to each allele on chromatin-associated or cytoplasmic RNA was compared using Qllelic (Mendelevich et al. 2021) or a two-sided binomial test, respectively. ***: p-value $< 0.001$ and proportion of allele 1 reads $< 0.4$ or $> 0.6$; **: p-value $< 0.01$ and proportion of allele 1 reads $< 0.45$ or $> 0.55$; *: p-value $< 0.05$ and proportion of allele 1 reads $< 0.45$ or $> 0.55$.
